# BOTTOM-UP EFFECTS AND DENSITY DEPENDENCE DRIVE THE DYNAMIC OF AN ANTARCTIC SEABIRD PREDATOR-PREY SYSTEM

**DOI:** 10.1101/2023.06.06.543885

**Authors:** Lise Viollat, Maud Quéroué, Karine Delord, Olivier Gimenez, Christophe Barbraud

## Abstract

Understanding how populations respond to variability in environmental conditions and interspecific interactions is one of the biggest challenges of population ecology, particularly in the context of global change. Although several studies have investigated population responses to climate change, very few have explicitly integrated interspecific relationships when studying these responses. Here, we aim to understand the combined effects of inter- and intraspecific interactions and environmental conditions on the demographic parameters of a prey-predator system of three sympatric seabird populations breeding in Antarctica: the south polar skua (Catharacta maccormicki), and its two main preys during the breeding season, the Adélie penguin (Pygoscelis adeliae) and the emperor penguin (Aptenodytes forsteri). We built a two-species integrated population model (IPM) with 31 years of capture-recapture and count data, and provided a framework that allows estimating demographic parameters and abundance of a predator-prey system in a context where capture-recapture data are not available for one species. Our results showed that predator–prey interactions and local environmental conditions affect differently south polar skuas depending on their breeding state of the previous year. Concerning prey-predator relationships, the number of Adélie penguin breeding pairs showed a positive effect on south polar skua survival and breeding probability, and the number of emperor penguin dead chicks showed a positive effect on the breeding success of south polar skuas. In contrast, there was no evidence for an effect of the number of south polar skuas on the demography of Adélie penguins. We also found an important impact of sea ice conditions on both the dynamics of south polar skuas and Adélie penguins. Our results suggest that this prey-predator system is mostly driven by bottom-up processes and local environmental conditions.

## INTRODUCTION

Marine ecosystems are difficult to study and quantify due to the large number of species involved, their complex interactions (interspecific and intraspecific), the diversity of environmental factors involved, and intricate mechanisms that interact together (Godfray & May, 2014). Fluctuations and changes in environmental conditions could interact with the different processes driving marine ecosystems (Stenseth et al., 2002), and even more in polar marine ecosystems, which are highly sensitive to global change (Smetacek & Nicol, 2005; Schofield et al., 2010; Doney et al., 2012). Seabirds are great indicators species to study the state of marine ecosystems and the impact of global changes (Regehr & Montevecchi, 1997; Cook et al., 2014; Velarde et al., 2019). As upper trophic-level long-lived predators, their dynamics reflect the impacts of climate change on lower trophic levels which are often difficult to assess (Barbraud & Weimerskirch, 2001; Jenouvrier et al., 2003; Piatt et al., 2007). Easier to study over the long term than other upper trophic-level marine predators, highly visible and nesting in large colonies in generally constant locations, they allow efficient data gathering and can be used as a proxy to better understand some ecologic processes on a large spatial and temporal scale (Stenseth et al. 2002; Hazen et al., 2019; Velarde et al., 2019). Moreover, numerous studies have showed that bottom-up (Cairns, 1987; Oro et al., 2007) and top-down processes (Oro & Furness, 2002; Hipfner et al., 2012) affect simultaneously and differently the demographic parameters of seabird populations (Votier et al., 2008; Horswill et al., 2014; 2016; Sauser et al., 2021). Thus, understanding the roles and importance of intrinsic and extrinsic factors on the dynamic of seabird populations is crucial in the context of global change (Horswill et al., 2016; Pacoureau, 2018). Quantifying the relative importance of environmental conditions, density dependence and interspecific relationships, such as prey-predator effects, on the dynamics of seabird populations could help us to better apprehend the relative importance of bottom-up and top-down processes on marine ecosystems (Quéroué et al., 2021) and better predict the future of species.

However, in order to better understand the dynamics of communities such as prey-predator systems, we need to consider not only the abundance of populations but also to follow a multi-trait approach studying the survival, recruitment, and reproductive parameters (Cook et al., 2014; Horswill et al., 2016; Perkins et al., 2018), as well as the structure of the populations including the different ages and states of individuals (Miller & Rudolf, 2011). Species responses to fluctuations of extrinsic factors like climatic conditions or food availability also differ depending on the species (Grosbois et al., 2008; Jenouvrier, 2013) or on the individuals ontogenic stage (Miller & Rudolf, 2011; Gervasi et al., 2015). This could lead to modifications of the population structure, of species interactions, or of the diversity within communities (Thomas et al., 2004). Assessing the responses to environmental fluctuations species by species could overlook the role played by the different interspecific interactions. Furthermore, intrinsic factors such as density-dependence effects or individual characteristics (ontogenic stage, age, size) could also influence the structure and dynamics of populations (Ashmole, 1963; Kramer et al., 2009).

Integrated population models (IPM) are a powerful framework that allows to estimate the state-dependent demographic parameters and the abundance of individuals within populations using different sources of information (count data, capture-recapture data, etc.) in a single analysis, simultaneously considering the various sources of uncertainty related to each type of data (Besbeas et al., 2002; Schaub & Abadi, 2011; Zipkin & Saunders, 2018). Recently, IPMs have been extended to multiple species for competition/parasitism (Péron & Koons, 2012) or for predator-prey interactions (Barraquand & Gimenez, 2019). Quéroué et al. (2021) developed a multi-specific IPM to assess the dynamics of a seabird predator-prey system using capture-recapture data for a predator (the brown skua *Catharacta lönnbergi*) and its main prey (the blue petrel *Halobaena caerulea*). However, obtaining capture-recapture data requires a substantial sampling effort and is not always possible to achieve for all species involved in predator-prey systems due to logistical or technical reasons. Thus, needed are multi-specific IPM for cases where capture-recapture and count data are only available for one species of the predator-prey system, the data concerning the other species being only time series of counts.

The aim of our study was to assess the effect of inter and intraspecific relationships, as well as environmental conditions on the demographic parameters within a predator-prey system of seabirds in Antarctica. To this end we developed a two species IPM combining 31 years of capture-mark-recapture (CMR) and count data for the south polar skua (*Catharacta maccormicki*) and count data for one of its main prey during the breeding season, the Adélie penguin (*Pygoscelis adeliae*). We also include, as a covariate into the model, the emperor penguin dead chicks (*Aptenodytes forsteri*), another major prey of the south polar skua.

The main strengths of our approach are that i) it efficiently combines multiple and different sources of data within a unified framework, ii) it allows the estimation of demographic parameters of a predator-prey system while assessing the impact of intrinsic and extrinsic factors on these demographic parameters in order to better understand the contribution of predator–prey interactions to population dynamics.

## MATERIALS & METHODS

### STUDY SITE AND SPECIES

We studied south polar skuas, Adélie penguins and emperor penguins at the Pointe Géologie archipelago, Terre Adélie, near the Dumont d’Urville French Research station (66°40’ S, 140°01’ E). Skuas and Adélie penguins nest during the austral summer (October-February) and emperor penguins breed during the austral winter (April-December). In this study, we focused on the skuas and Adélie penguins breeding on the major islands of the archipelago (Le Mauguen, Pétrels, Rostand, Bernard, Lamarck, and Nunatak), representing 90% and 78% of the total populations of the archipelago for both species, respectively. Changes in the population sizes of the two species during the study period are shown in Figure 1.

**FIGURE 1:**
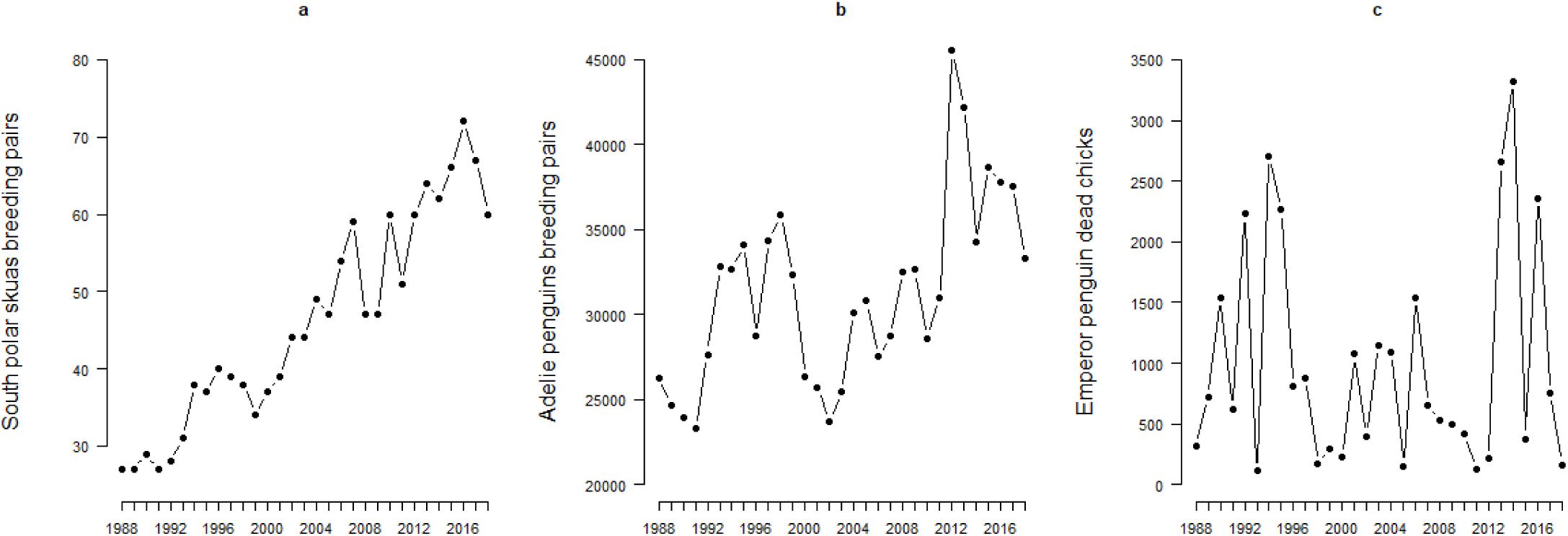
Time series showing the variation of the number of south polar skua breeding pairs (a), number of Adélie penguin breeding pairs (b) and emperor penguin dead chicks (c) from 1988 to 2018

South polar skuas are medium size seabirds breeding all around the coast of Antarctica. They have various and opportunistic diets as well as different foraging technics depending on the breeding localities and seasons. Strongly territorial during the breeding season, skuas are faithful to their breeding sites and to their partner (Eklund, 1961; Jouventin & Guillotin, 1979; Ainley et al., 1990). Breeding pairs form in late October, generally two eggs are laid in November and hatch around mid-December (Barbraud & Weimerskirch 2006). Chicks fledge about 50 days later. The parents feed their chicks until their departure of the breeding sites, around the end of March (Le Morvan et al., 1967). At the end of the breeding season, skuas from the Pointe Géologie archipelago migrate towards the east coast of Japan (35-45°N) to wintering (Weimerskirch et al., 2015). Juveniles will come back at Pointe Géologie 4 to 7 years after their departure to breed for the first time (Pacoureau et al., 2019). At Pointe-Géologie during the breeding season south polar skuas feed almost exclusively on penguins’ chicks and eggs, as scavengers or as predators. The breeding cycle of skuas is synchronized with the one of penguins (Table 1). From their arrival at Pointe Géologie until the sea ice breaks up in December or January, skuas feed on dead chicks of emperor penguins (Pryor, 1968). From November until the end of the breeding season they mainly feed on the eggs and chicks of Adélie penguins (Jouventin & Guillotin, 1979; Ainley et al., 1990).

**TABLE 1:**
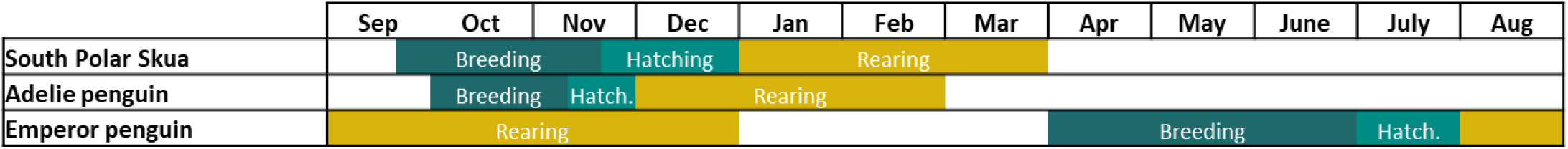
Reproductive phenology of the south polar skua and its two main preys: the emperor penguin and the Adélie penguin during the summer period on the Pointe Géologie archipelago (Pacoureau, 2018; Jouventin et al., 1995). The period of breeding includes pair formation, laying and incubation of the eggs, and the period of rearing corresponds to the time when parents care for their chicks until they fledge. The blank periods correspond to wintering and migration.

Adélie penguins are medium size penguins living all around the Antarctic coast. During the breeding season, Adélie penguins form dense colonies on rocky ridges on the Antarctic coast or on islands. They arrive at the colonies in October (Barbraud & Weimerskirch 2006). Females lay in general two eggs, which will hatch 30 days later. From hatching, the parents make numerous trips between the colony and the pack ice to forage food for the chicks (Widmann et al. 2015). Highly dependent on sea ice conditions for their food, they feed mainly on crustaceans (in particular different species of krill including the Antarctic krill *Euphausia superba*, Cherel 2008). All chicks fledge at the end of February (Marchant & Higgins, 1990; Ainley, 2002). Outside the breeding period, Adélie penguins from Pointe Géologie migrate towards the northwest, pursuing the pack-ice edge within an area spanning around the colony of 1 900 000 km^2^ (Thiebot et al., 2019).

Emperor penguins are large size penguins living all around the Antarctic coast and are the only birds breeding directly on sea ice during the austral winter (Jouventin et al., 1995). There are highly dependent on sea ice for their food (fish, crustacean, cephalopods) and as a breeding platform (Barbraud et al., 2011; Jenouvrier et al., 2014; 2019). Each breeding season, they form colonies of thousands of individuals (Fretwell et al., 2014). Females lay a single egg, which will be carefully protected between the feet of the male until hatching in July (Table 1). From hatching, the parents make numerous trips between the colony and the sea ice edge to forage food for the chick (Zimmer et al. 2008). Survival of emperor penguin chicks during the breeding season is highly sensitive to the sea ice conditions between September and November and is negatively affected by longer distances between colony and fast ice edge (Labrousse et al., 2021). In December, all adults leave the colony, shortly followed by the chicks. Four to six years later, juveniles reach their sexual maturity and come back to breed (Mougin & van Beveren 1979). Outside the breeding season, emperor penguins live exclusively within the pack ice where they feed on fishes, crustaceans (krill), and cephalopods (Borboroglu & Boersma, 2015).

### COUNT AND CAPTURE-RECAPTURE DATA

We used demographic data for both skuas and Adélie penguins collected during the breeding season from 1988/1989 to 2018/2019 (thereafter named 1988 to 2018). For emperor penguins, we used a time series data providing information on the annual number of dead chicks which was used as a covariate in the IPM. For skuas, three types of data were used: count data corresponding to the number of breeding pairs (Ys), capture-recapture (CR) data of adult individuals observed in the monitored area and the number of immigrants, i.e. new individuals observed for the first time (not ringed) at the colony (N_im_). Every year, the locations of the different nests were spotted during the laying period. The monitored area occupied being relatively small (about 80 ha), with no vegetation, and given the conspicuous behaviour of the breeding skuas when defending their territories, we assumed that all active nests were detected each year (Pacoureau et al., 2018). A major part of the population is marked using stainless steel and plastic bands engraved with a unique alphanumeric code. Every year during the breeding season, from mid-October to mid-April, nesting territories, as well as a zone where nonbreeding individuals are known to roost, were visited every two weeks in average to determine the breeding status of each bird (Pacoureau et al., 2019).

The breeding status of each individual was defined as:

- Breeder (B): individual that is being seen on a nest with at least one egg
- Nonbreeder (NB): individual never seen on a nest with eggs or chicks
- Failed breeder (FB): breeding individual whose eggs did not hatch and/or chicks did not survive until

fledging

- Successful breeder (SB1): breeding individual with one chick fledged
- Successful breeder with 2 chicks (SB2): breeding individual with two chicks fledged

We also defined an uncertain state (C) for individuals which could not be assigned with certainty to one of the five breeding states described above. As juveniles come back at the colony when they became adults and in age to breed (from 4 years or more), we did not have observation data for them before they came back at Pointe Géologie. The annual number of breeding pairs of skuas, i.e. count data (Y_S_), was estimated as the number of nests identified and occupied by breeding individuals (Pacoureau et al., 2018).

For Adélie penguins, two types of count data were used: the number of breeding pairs (Y_A_) and the number of chicks ready to fledge (Y_P_). Breeding pairs are visually counted every year, between 15 and 18 December, during the laying period. Live chicks ready to fledge are counted before their departure at sea, between 3 and 6 February. See Barbraud et al. (2020) for more information on monitoring protocols.

### INTEGRATED POPULATION MODEL

We built a two species IPM that combines count and capture-recapture data that allows estimating abundance and demographic rates of south polar skuas and Adélie penguins. More specifically, we connected one IPM for the skuas and a state-space model for Adélie penguins through explicit predator-prey relationships (Barraquand and Gimenez 2019). We incorporated the effects of species-specific demographic parameters such as survival and breeding parameters. The two models used are structured according to life-history traits (Figure 2). We used Poisson (Po) and binomial (Bin) distributions to account for demographic stochasticity. The details of the model are presented in Appendix 1.

**FIGURE 2:**
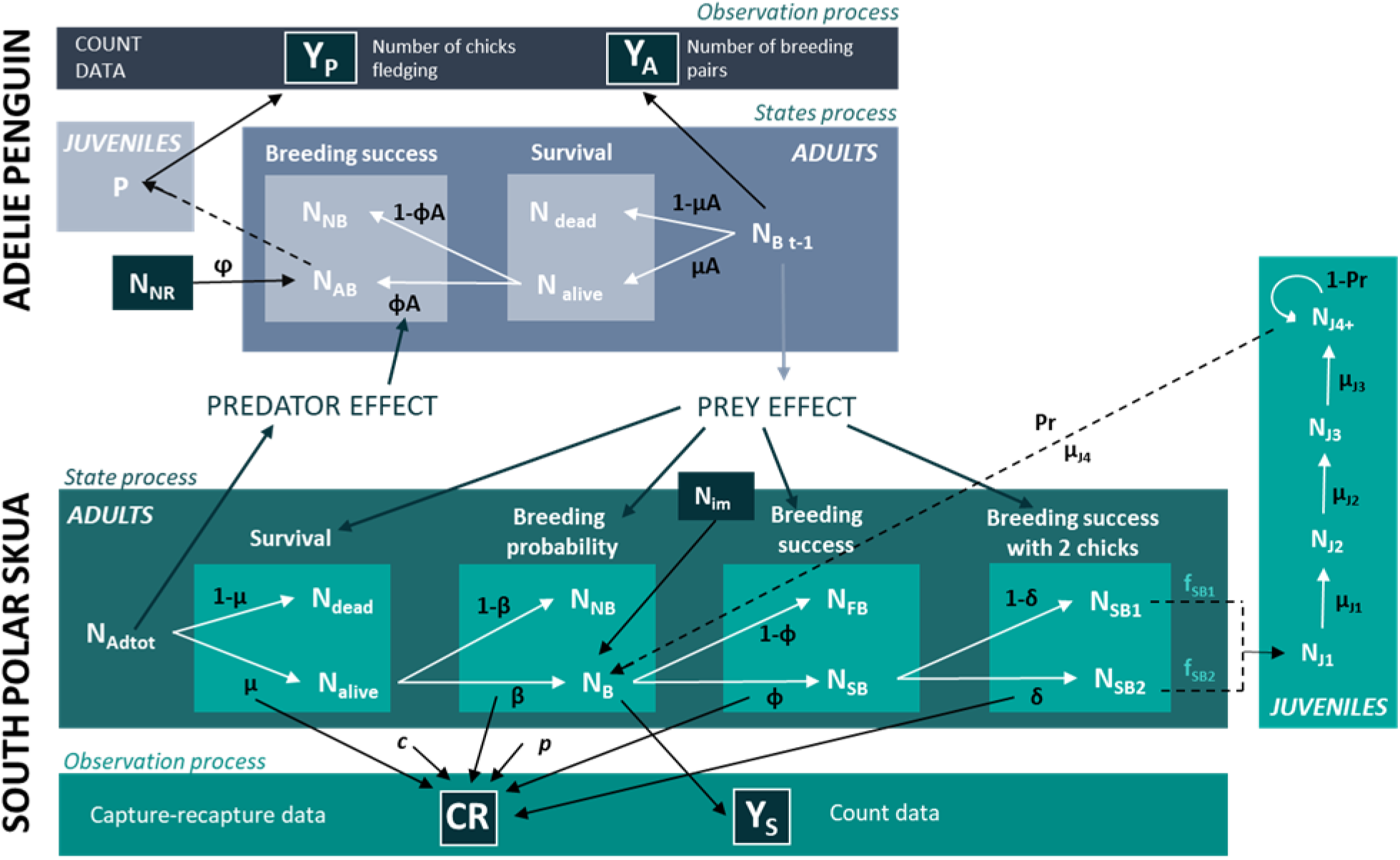
Structure of the multispecies integrated population model for south polar skuas and Adélie penguins. Writing in white indicates the different states, and in black the demographic parameters. Four types of data were used: capture-recapture data (CR), counts of breeding pairs of skuas (Y_S_), counts of breeding pairs of Adélie penguins (Y_A_) and the number of Adélie penguin chicks just before fledging (Y_P_). For skuas, the parameters estimated by the model were the apparent adult survival (μ), the apparent juvenile survival (μ_J1_ to μ_J4_), the probability of first breeding (Pr), the breeding probability (β), the breeding success (φ), the breeding success with 2 chicks (δ), the detection probability (p) and the probability to assign an individual to its real state (c). The skua fecundity parameters (f_SB1_ and f_SB2_) were fixed according to the breeding states (with one chick: SB1 or two chicks: SB2). For Adélie penguins, the parameters estimated by the model were the adult apparent survival (μ_A_) and the breeding success (φ_A_). The Adélie penguin recruitment rate (ϕ) was fixed. For skuas, the state variables estimated by the model were the total number of adults (N_adtot_), dead (N_dead_), alive (N_alive_), breeders (N_B_), non-breeders (N_NB_), failed breeders (N_FB_), successful breeders with one chick (N_SB1_), successful breeders with 2 chicks (N_SB2_), the number of juveniles of 1 to 4 years old or older (N_J1_ to N_J4+_). The number of immigrants, i.e. new individuals observed for the first time (not ringed) at the colony (N_im_) was a fixed vector. For the Adélie penguin, the state variables estimated by the model were the number of breeding pairs (N_AB_), alive (N _alive_), dead (N_dead_), adult non-breeders (N_NB_), fledged chicks (P). N_NR_ corresponds to the number of Adélie penguin new breeders (local recruitment and immigration).

Demographic parameters of south polar skuas and Adélie penguins could be affected by different covariates including interspecific predator-prey relationships, intraspecific density-dependence, and environmental covariates such as temperature or sea ice conditions. To estimate those effects, we used logit linear regressions on the estimation of the demographic parameter within the IPM. An example is shown Appendix 2.

All covariates were standardized to compare the relative contribution of the effects. We calculated the 95% credible intervals (CRI) for the regression coefficients α, and the probabilities of having a negative slope value (PN) or of having a positive slope value (PP). A regression coefficient was considered significant when its probability of being negative (PN) was greater than 90% or less than 10% (equivalent to a probability of being positive (PP) at 90%).

#### Model implementation

Bayesian posterior distributions of the multispecies IPM were approximated with Markov chain Monte Carlo (MCMC) algorithms. Two independent chains MCMC of 30 000 iterations were used, with a burn-in period of 10 000 iterations. Gelman-Rubin convergence diagnostics (Brooks & Gelman, 1998) were below 1.1 for each parameter and the mixing of the chains was satisfactory. The analyses were performed using JAGS (Plummer, 2014; version 4.3.0) and program R (R Core Team, 2020; R version 4.0.5).

### COVARIATES AND HYPOTHESES

In the following, we detail the covariates used and how we hypothesize they may affect the dynamics of skuas and Adélie penguins. The covariates tested on the different demographic parameters for the two species are resumed in Table 2.

**TABLE 2:** Summary of the covariates tested on the demographic parameters of south polar skuas (breeders B and non-breeders NB) and Adélie penguins, and time periods considered for each demographic parameter.

| PARAMETERS | COVARIATES TESTED | TIME PERIOD |
| --- | --- | --- |
| **Skuas (B and NB)** |  |  |
| Adult apparent survival $\mu$ | PP, DD, SIC, AT, SSTa | Spring, summer |
| Breeding probability $\beta$ | PP, DD, SIC, AT, SSTa | Spring |
| Breeding success $\phi$ | PP, DD, SIC, AT | Spring, summer |
| Breeding success with 2 chicks $\delta$ | PP, DD, SIC, AT | Spring, summer |
| **Adelie penguins** |  |  |
| Breeding success $\phi_A$ | PP, DD, SIC | Winter, spring, summer |

*Notation:* PP, predator-prey interactions, i.e. number of Adélie penguin breeding pairs and emperor penguin dead chicks for skua demographic parameters, and number of skua breeding pairs for Adélie penguin demographic parameters; DD, intraspecific density dependence; SIC, sea ice concentration; AT, air temperature; SSTa, sea surface temperature anomalies

#### Predator-prey interactions

To estimate the effect of interspecific predator-prey relationships between skuas and Adélie penguins, i.e. the dependence of demographic parameters of one species according to the abundance of another species, we used the following state variables: the number of adult (breeders and non-breeders) skuas (N_adtot_) and the number of Adélie penguin breeding pairs (N_AB_) estimated by the multispecies IPM. We also used the number of dead emperor penguin chicks as a covariate to test its effect on the skua demographic parameters. Carcasses of dead emperor penguin chicks are counted every day from the end of July to December until the sea ice breaks up. We hypothesized that a high availability of preys (number of Adélie penguin breeding pairs and dead emperor penguin chicks) will favour the survival, breeding probability and breeding success of south polar skuas (Newton, 1998). As the carcasses of dead emperor penguin chicks are available at the beginning of the skua breeding season, we supposed that an important number of dead chicks could favour the breeding probability (β) of skuas. An important number of breeding Adélie penguins and emperor penguin dead chicks would ensure their accessibility to food resources during the breeding season, and thus would increase their survival (μ) as well as the breeding successes (φ and δ). On the opposite, the predation on eggs and chicks of Adélie penguins by south polar skuas should have a negative effect on the breeding success (φ_A_) of Adélie penguins (Horswill et al., 2016).

#### Density dependence

We estimated intraspecific relationships for both species using the number of adults (breeders and non-breeders) skuas (N_adtot_) and the number of Adélie penguin breeding pairs (N_AB_) estimated by the IPM. We supposed that an important number of individuals of the same species within the colony could have a negative effect on the survival and breeding success, through an increased intraspecific predation on eggs and chicks (Furness, 1987) and competition for food resources and/or for high quality nesting sites (Hunt et al., 1986; Dhondt et al., 1992; Turchin et al., 1995; Newton, 1998). At high densities, skuas and Adélie penguins would be less efficient to raise their chicks and thus would have lower breeding success (φ, δ) than at low densities.

#### Environmental covariates

We considered several covariates suspected to affect populations of skuas and Adélie penguins, directly and through perturbations of the food web (Barbraud & Weimerskirch, 2001; Jenouvrier et al., 2003). For both Adélie penguins and south polar skuas, we supposed that local climatic conditions encountered by the parents and their offspring during the breeding season, as well as the climatic conditions encountered outside the breeding season by carry-over effects, will play a determining role in the survival, breeding probability and breeding success by affecting food availability and energy expenditure. Temporal variations for the environmental covariates considered over the study period (1988-2018) are represented in Figure 3.

**FIGURE 3:**
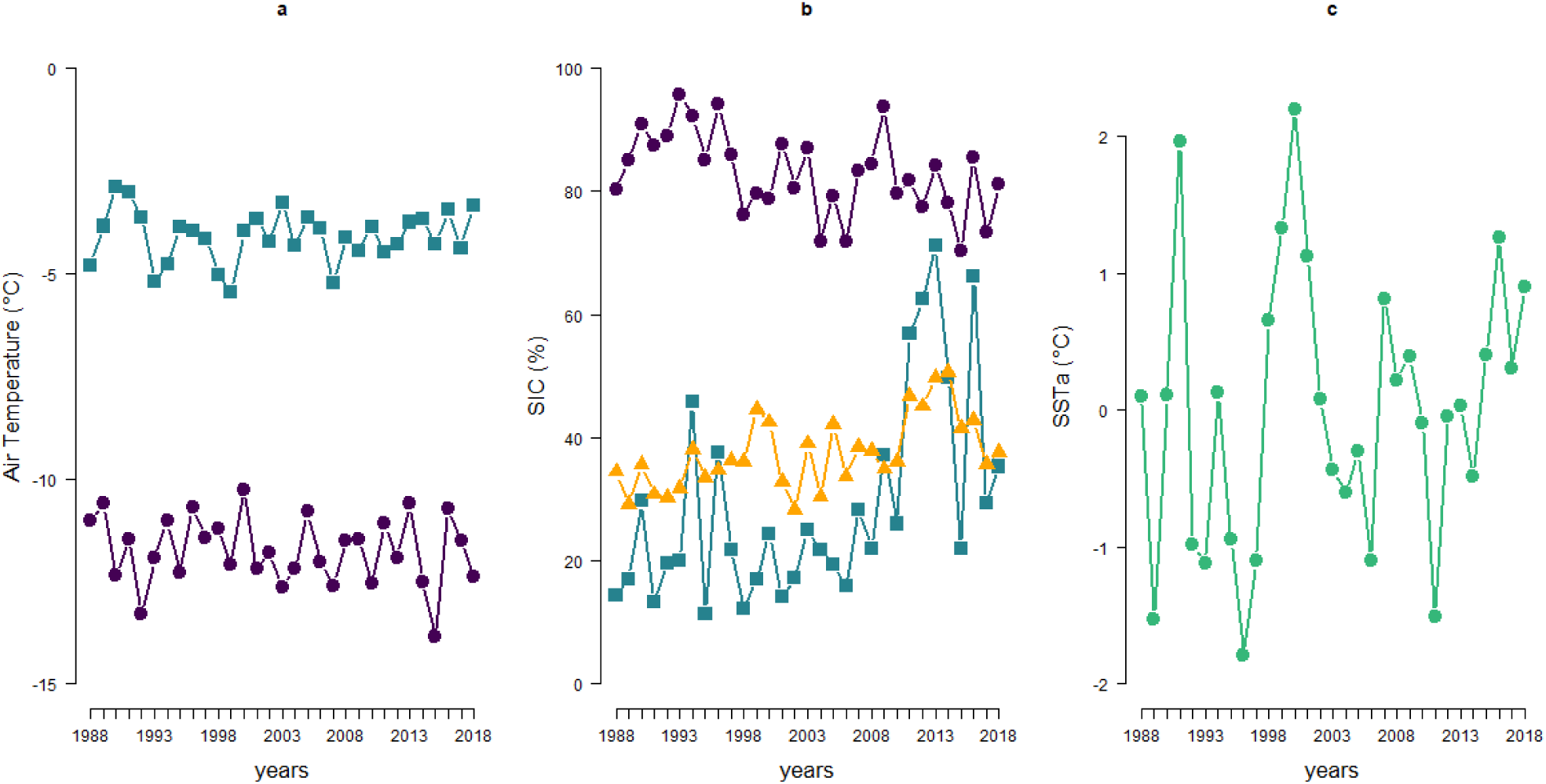
Time series showing the variation of the environmental covariates used in the model from 1988 to 2018. (a) Average air temperature recorded at Dumont d’Urville station in spring (purple circle) and summer (blue square). (b) Average sea ice concentration (SIC) in spring (purple circle), summer (blue square) and winter (orange triangle). (c) Average sea surface temperature anomaly (SSTa) between May and June in the wintering area of skuas.

#### Air temperature (AT)

It was measured daily at the Dumont d’Urville (DDU) station, localized on Ile des Pétrels, Pointe Géologie archipelago. Data were downloaded from British Antarctic Survey (https://legacy.bas.ac.uk/met/READER/data.html). Temperature data were averaged in two periods: spring (September-November) and summer (December-March). The first period corresponds to the breeding period when skuas arrive at the colonies, make courtship rituals, form pairs, prepare the nest, lay, and incubate the eggs. The second period corresponds to chick rearing from hatching until fledging. We supposed a negative effect of low air temperatures on demographic parameters (i.e. survival, breeding probability and breeding success) of skuas. Low temperatures could cause an extra energetic cost due to thermoregulation and water loss, which could penalize parents, lead to smaller eggs laid, and to a higher mortality of chicks (Spellerberg, 1969; Emmerson & Southwell, 2011).

#### Sea ice concentration (SIC)

Monthly SIC data were averaged into three periods: late spring (September–November), summer (December–March), and winter (April-August). For spring and summer data, we considered the area around DDU station (65°50’-66°50’ N, 139°50’-140°50’ E) corresponding to the area where emperor penguins, Adélie penguins, and skuas forage during their breeding season (Pacoureau et al., 2018). The winter data area was extended at (60°00’-66°50’ N, 100°00’-150°00’ E), corresponding to the foraging zone of Adélie penguins during their wintering (Thiebot et al., 2019). The data came from the National Oceanic and Atmospheric Administration (http://iridl.ldeo.columbia.edu/SOURCES/.NOAA/.NCEP/.EMC/.CMB/.GLOBAL/.Reyn_SmithOIv2/.monthly/.sea_ice/index.html). As the abundance and distribution of their main prey (Antarctic krill or ice krill *E. crystallorophias*) depend entirely on sea ice (Ainley, 2002; Croxall et al., 2002; Forcada et al., 2006; Flores et al., 2012), Adélie penguins are highly sensitive to changes in oceanic conditions, and even more during the breeding season when the foraging area is limited (Weimerskirch, 2007). Numerous studies showed a link between sea ice conditions near the colony, as a proxy of krill availability, and the breeding success of Adélie penguins (Marchant & Higgins, 1990; Ainley, 2002; Emmerson & Southwell, 2008; Emmerson et al., 2011; Barbraud et al., 2015; Barreau et al., 2019). We supposed that high SIC during wintering and at the beginning of the breeding season (spring) will allow Adélie penguins to develop a better body condition and will reduce the cost of reproduction and so favour their breeding success (Emmerson & Southwell, 2011; Grilli et al., 2018; Dunn et al., 2020). On the opposite, a high SIC during summer will increase foraging trips lengths and energetic costs for Adélie penguins raising chicks (Emmerson & Southwell, 2008), which could lead to a decrease in breeding success (Ropert-Coudert et al., 2014; Barbraud et al., 2015).

We thus hypothesised that sea ice conditions will affect the breeding success of Adélie penguins, as well as the south polar skua dynamics through bottom-up processes (Frederiksen et al., 2006). High SIC conditions in spring could favour the breeding probability and breeding success of skuas, as there would be more potential food resources available for the predator. Likewise, high SIC conditions in summer may increase the survival and breeding success of skuas, due to the greater availability of penguin chick carcasses during the rearing period of their chicks.

#### Sea surface temperature anomalies (SSTa)

SSTa were used as a proxy of food availability in the wintering zone of south polar skuas, i.e. east of the Japan sea (32°50′-44°50′ N, 139°50′-149°50′E). Data were averaged between May and June and came from the National Oceanic and Atmospheric Administration. High SSTa generally reflects poor local environmental conditions for zooplankton development, and so potentially a lower availability of food for skuas, by repercussion along the food web. (Frederiksen et al., 2006; Barbraud et al., 2012; Weimerskirch et al. 2015; Hazen et al., 2019). We supposed a negative effect of high SSTa during the wintering on the survival of skuas (μ), but also on their breeding parameters through carry-over effects, i.e. events encountered by an individual before the breeding season which will influence its breeding performance during the following season (Norris, 2005; Harrison et al., 2011; O’Connor et al., 2014).

Responses to covariates were modelled as dependent on the breeding state at the previous breeding season. In our IPM, we assessed the different demographic parameters (µ, β, φ, δ) of skuas according to their breeding state during the previous breeding season (breeder thereafter noted B, or non-breeder thereafter noted NB). Non-breeders were individuals that reached sexual maturity but do not reproduce (Penteriani et al., 2011) or skipped breeding due to a poor body condition or because they could not find a partner or a territory to breed (Ashmole, 1963). We supposed different values of the demographic parameters of skuas depending on their breeding state on the previous year, as well as the different responses to environmental factors and intra and inter-specific relationships (Ashmole, 1963).

## RESULTS

The number of south polar skua breeders and the number of Adélie penguin breeding pairs obtained from model estimates and observed are shown in Figure 4.

**FIGURE 4:**
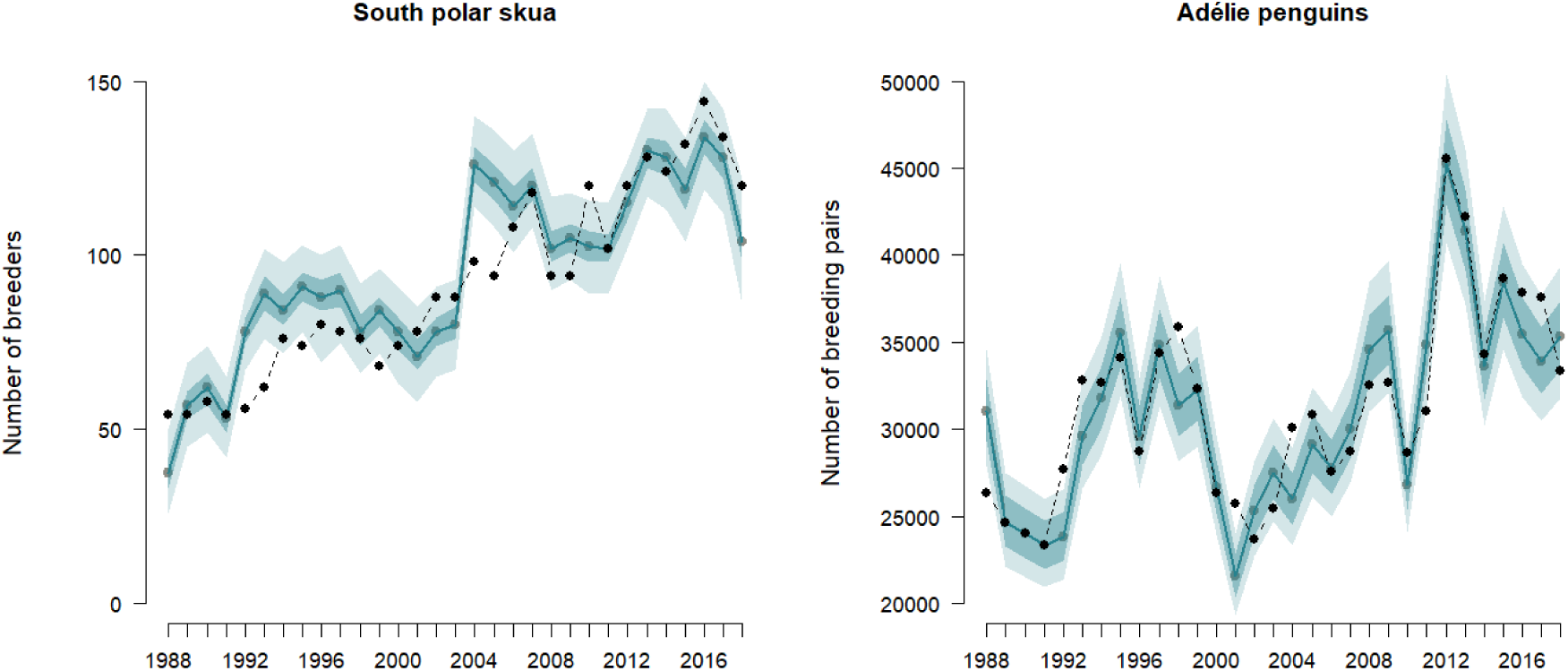
Estimated number of south polar skua breeders and Adélie penguin breeding pairs at Pointe Géologie from 1988 to 2018. Solid blue line and blue dots represent the means of marginal posterior distributions. Shaded areas are the 50% and 95% credibility intervals. Black dots and dashed line represent the observed number of breeding skuas and penguins (count data).

### DEMOGRAPHIC PARAMETERS

Mean values of demographic parameters for skuas and Adélie penguins estimated by the model are presented in Table 3, and their values for each year are shown in Figure 5. Estimates of capture probability of skuas are shown in Appendix 3. Variation of the estimated number of skuas for the different breeding states between 1998 and 2018 are shown in Appendix 4.

**FIGURE 5:**
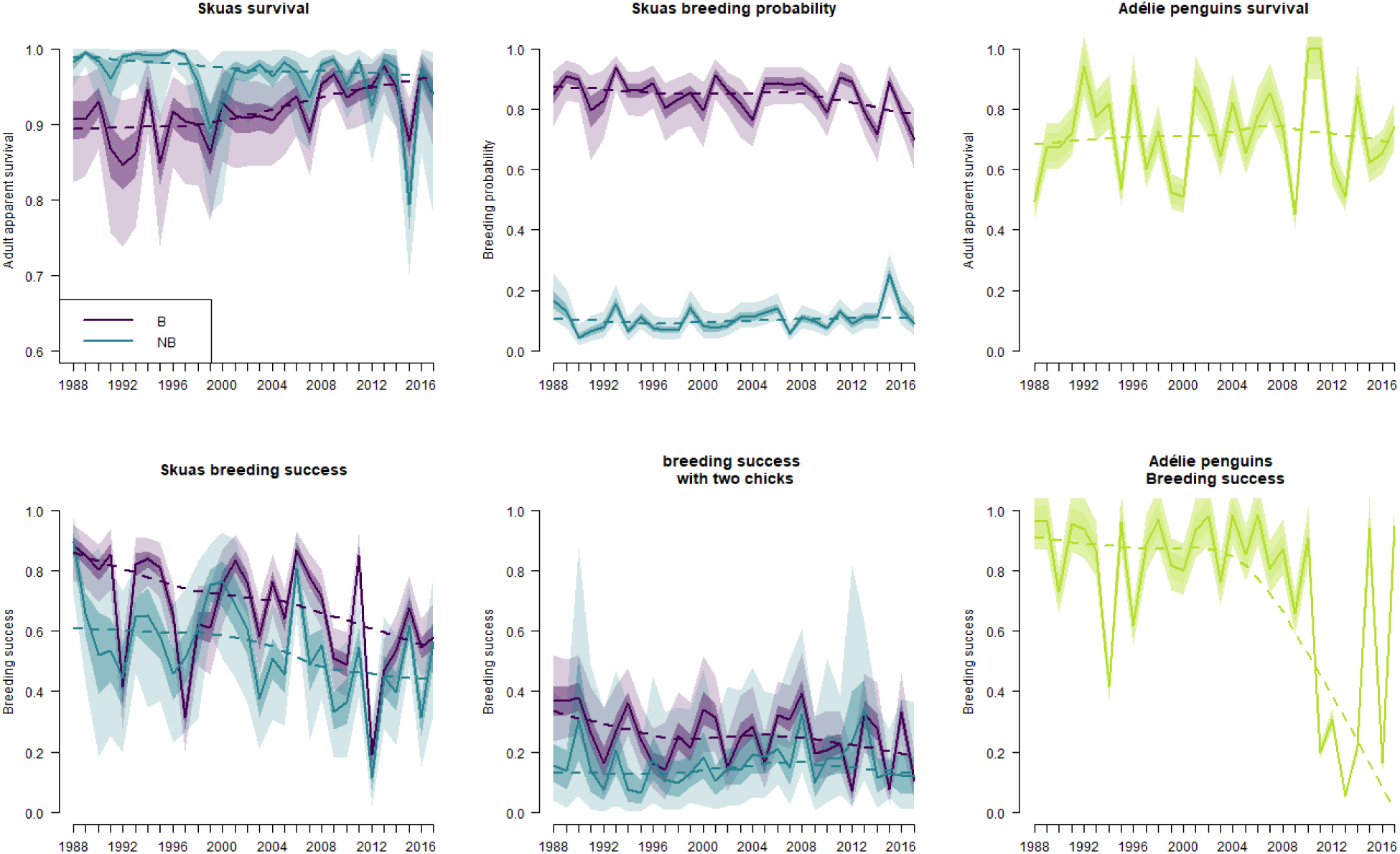
Estimated demographic parameters for south polar skuas by years, depending on their breeding state at the previous breeding season (B: breeder, in purple, and NB: non-breeder, in blue), and Adélie penguins (in green) of Pointe Géologie from 1988 to 2018. Solid lines represent the means of marginal posterior distributions and dashed lines the marginal posterior distributions smoothed by the lowess function in R. Shaded areas are the 50% and 95% credibility intervals.

**TABLE 3:** Mean demographic parameters estimates for south polar skuas depending on their breeding states (B: breeder and NB: non-breeder) during the previous breeding season and for Adélie penguins estimated by the model from 1988 to 2018 for the populations of Pointe Géologie. 95 % CRI: 95% credibility intervals

| PARAMETERS |  | Mean | 95% CRI |
| --- | --- | --- | --- |
| **South polar skuas** |  |  |  |
| Adult apparent survival $\mu$ | B | 0.914 | [0.847, 0.963] |
|  | NB | 0.958 | [0.892, 0.994] |
| Breeding probability $\beta$ | B | 0.842 | [0.753, 0.913] |
|  | NB | 0.107 | [0.065, 0.159] |
| Breeding success $\phi$ | B | 0.662 | [0.544, 0.77] |
|  | NB | 0.532 | [0.323, 0.736] |
| Breeding success with two chicks $\delta$ | B | 0.251 | [0.145, 0.383] |
|  | NB | 0.182 | [0.032, 0.446] |
| **Adélie penguins** |  |  |  |
| Adult apparent survival $\mu_A$ | | 0.724 | [0.685, 0.764] |
| Breeding success $\phi_A$ | | 0.751 | [0.748, 0.754] |

The mean apparent adult survival (μ) for skuas was 0.936 ± 0.029, with a lower survival of 4.4% for B skuas than for NB skuas (Figure 5). Survival of the two breeding states showed similar inter-annual variations. However, the survival of B skuas increased by 3.3% during the study period while the survival for NB skuas decreased by 4%. Adélie penguin apparent survival (μ_A_) was 0.724 ± 0.023. The breeding probability (β) for B skuas was 73% higher than the breeding probability for NB skuas. The breeding success (φ) for B skuas was 12.9% higher than for NB skuas, and the breeding success with two chicks (δ) was 6.9% higher for B skuas than for NB skuas. The two breeding success parameters for both breeding states showed similar variations through time and decrease by 31.6% for breeding success and by 14.9% for breeding success with two chicks. Adélie penguin breeding success (φ_A_) was 0.751 ± 0.002.

### EFFECT OF ENVIRONMENTAL COVARIATES AND POPULATION DENSITIES

All results obtained when estimating the effects of the different environmental covariates and population densities on demographic parameters of skuas and Adélie penguins are shown in Appendix 5 and resumed in Table 4. Only the significant effects will be interpreted here.

**TABLE 4:**
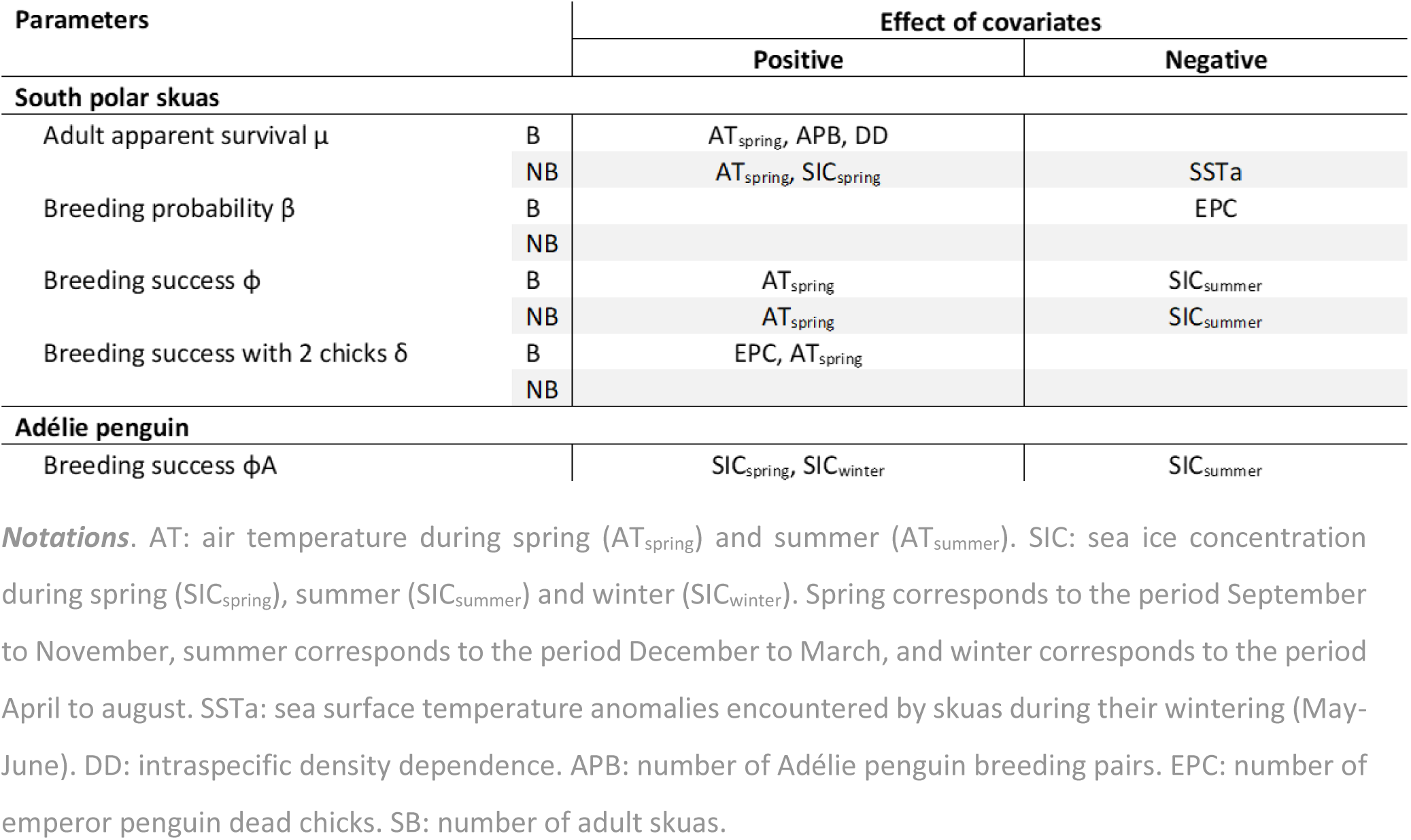
Estimates of the effects of the environmental covariates and population densities on the demographic parameters of south polar skuas depending on their breeding state at the previous breeding season (B: breeder, and NB: non-breeder), and of Adélie penguins. Only the significant effects (i.e. estimated effect with a probability of being negative greater than 90% or less than 10%) are shown.

| Parameters |  | Effect of covariates |  |
| --- | --- | --- | --- |
|  |  | Positive | Negative |
| **South polar skuas** |  |  |  |
| Adult apparent survival $\mu$ | B | AT<sub>spring</sub>, APB, DD | |
|  | NB | AT<sub>spring</sub>, SIC<sub>spring</sub> | SSTa |
| Breeding probability $\beta$ | B | | EPC |
|  | NB |  |  |
| Breeding success $\phi$ | B | AT<sub>spring</sub> | SIC<sub>summer</sub> |
|  | NB | AT<sub>spring</sub> | SIC<sub>summer</sub> |
| Breeding success with 2 chicks $\delta$ | B | EPC, AT<sub>spring</sub> | |
|  | NB |  |  |
| **Adélie penguin** |  |  |  |
| Breeding success $\phi_A$ | | SIC<sub>spring</sub>, SIC<sub>winter</sub> | SIC<sub>summer</sub> |

**Notations.** AT: air temperature during spring (AT<sub>spring</sub>) and summer (AT<sub>summer</sub>). SIC: sea ice concentration during spring (SIC<sub>spring</sub>), summer (SIC<sub>summer</sub>) and winter (SIC<sub>winter</sub>). Spring corresponds to the period September to November, summer corresponds to the period December to March, and winter corresponds to the period April to august. SSTa: sea surface temperature anomalies encountered by skuas during their wintering (May-June). DD: intraspecific density dependence. APB: number of Adélie penguin breeding pairs. EPC: number of emperor penguin dead chicks. SB: number of adult skuas.

#### Density-dependence

We estimated a positive density-dependence for the relationship between the apparent survival of B skuas (μ_B_) and the number of adult skuas (slope = 0.882, 95% CRI [0.687, 0.996]), as well as a positive density-dependence between the breeding success of Adélie penguin (φ_A_) and the number of Adélie penguin breeding pairs (slope = 0.999, 95% CRI [0.998, 1]).

#### Predator-prey relationships

We estimated several interspecific relationships between the number of prey (Adélie penguin breeding pairs and emperor penguins dead chicks) and the demographic parameters of the predator, the south polar skua (Figure 6). The apparent survival (μ_B_, slope = 0.76, 95% CRI [0.351, 0.993]) for B skuas increased with Adélie penguin breeding pairs. Even though the effects were less clear (95% CRI including zero), there was evidence for a negative relationship between the number of emperor penguin dead chicks and the breeding probability for B skuas (β_B_, slope = −0.206, 95% CRI [-0.458, 0.016]). On the other hand, we also detected that the probability of having two chicks (δ_B_, slope = 0.309, 95% CRI [-0.038, 0.695]) for B skuas was positively linked to the number of emperor penguin dead chicks. No significant relationship was found between the number of skuas and the breeding success of Adélie penguins. However, the number of adult skuas was positively correlated with the number of Adélie penguin breeding pairs (Pearson correlation index = 0.62).

**FIGURE 6:**
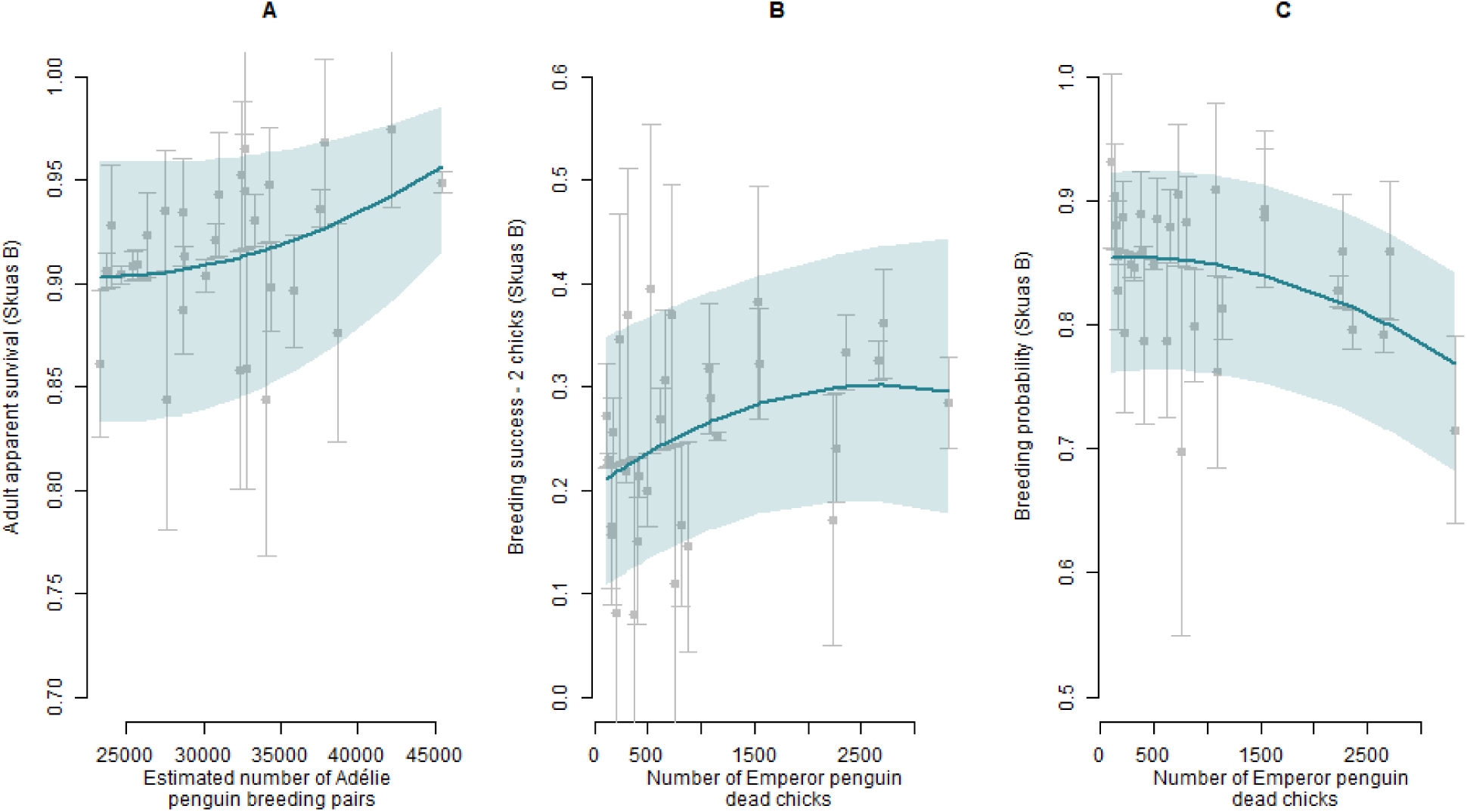
Interspecific relationships: effects of prey abundance on the demographic parameters of the south polar skua. A – Effect of the estimated number of Adélie penguin breeding pairs on survival of B skuas (µ_B_). B – Effect of the number emperor penguin dead chicks on breeding success with two chicks (δ_B_) of B skuas. C – Effect of the number emperor penguin dead chicks on breeding probability (β_B_) of B skuas. Solid line represents the modeled relationship obtained from the covariate model and shaded areas represent the 95% credibility intervals. Points represent demographic parameter estimates each year plotted against covariate values, with the standard deviations as errors bars. Demographic parameters of south polar skuas depend on their breeding state at the previous breeding season (B: breeder, and NB: non-breeder).

#### Environmental factors

We found positive relationships between air temperature in spring and breeding success of skuas. The breeding success of B skuas (φ_B_, slope = 0.251, 95% CRI [-0.168, 0.615]) and NB skuas (φ_NB_, slope = 0.289, 95% CRI [-0.085, 0.691]), as well as the probability to raise two chicks for B skuas (δ_B_, slope = 0.262, 95% CRI [-0.047, 0.615]) were higher when air temperatures between September and November increased. Higher temperatures in spring increased apparent survival of B skuas (μ_B_, slope = 0.225, 95% CRI [-0.08, 0.539]) and of NB skuas (μ_NB_, slope = 0.435, 95% CRI [-0.131, 0.942]). We did not find any significant effect of the air temperature in summer on the apparent survival or breeding probability and success of skuas.

Higher SIC during the breeding season was correlated with several demographic parameters of south polar skuas and Adélie penguins (Figure 7). The breeding success of Adélie penguins (φ_A_) was negatively related with SIC in winter (slope = −0.981, 95% CRI [-0.999, −0.954]) and in spring (slope = −0.643, 95 % CRI [-0.663, - 0.618]). The breeding success of Adélie penguins (φ_A_, slope= −0.378, 95% CRI [-0.404, −0.353]), as well as the breeding success of both B skuas (φ_B_, slope = −0.462, 95 % CRI [-0.882, −0.044]) and NB skuas (φ_NB_, slope = - 0.525, 95% CRI [-0.916, −0.121]) decreased with higher SIC conditions in summer. We also found positive relationships between the apparent survival of NB skuas (μ_NB_, slope = 0.482, 95% CRI [-0.199, 0.956]) and higher SIC in spring. On the other hand, when NB skuas encountered higher SSTa in their wintering area (May-June), their apparent survival was lower (μ_NB_, slope = −0.544, 95% CRI [-0.972, 0.251]).

**FIGURE 7:**
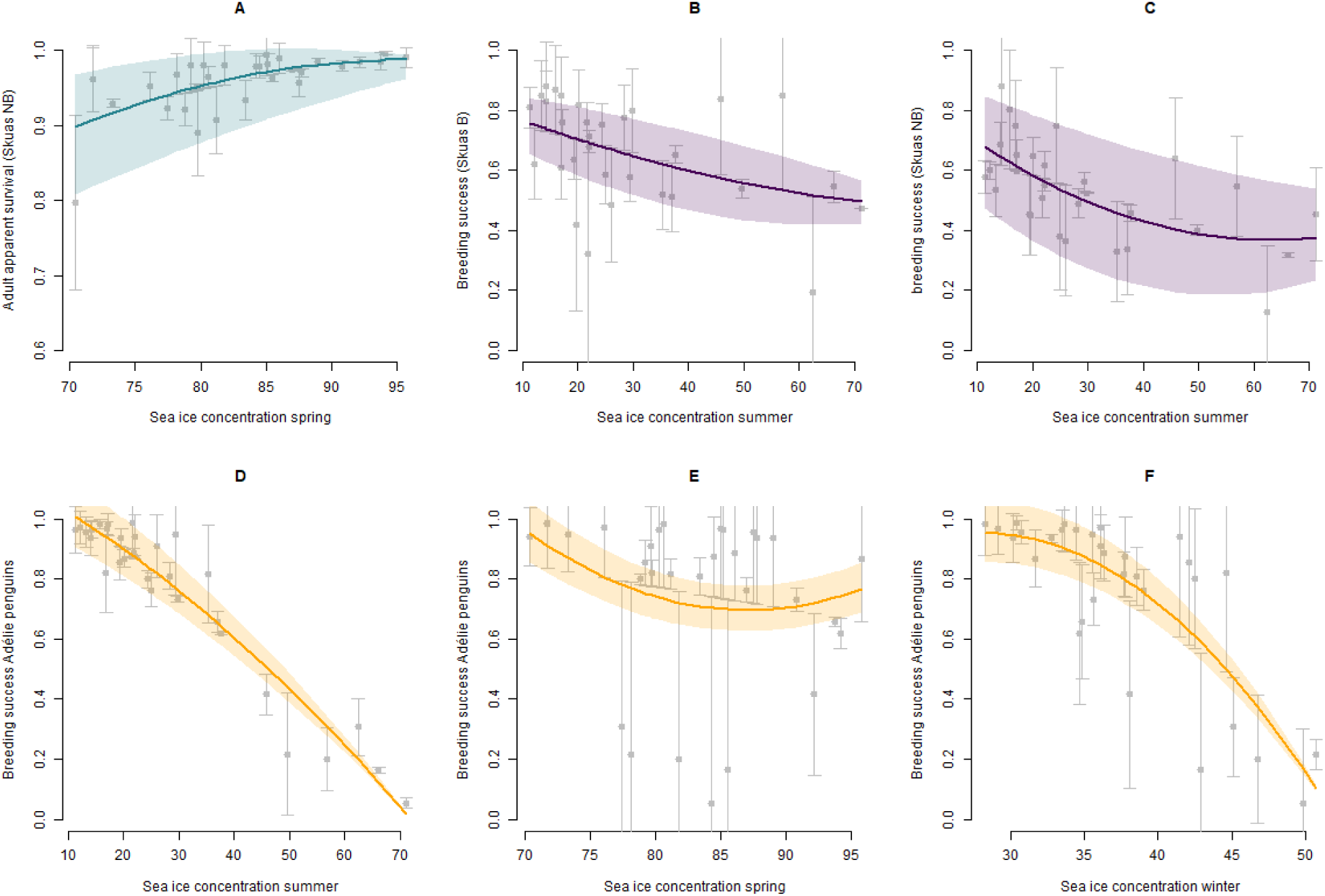
Effects of sea ice concentration (SIC). Effect of SIC during austral spring (September-November) on (A) adult apparent survival of NB Skuas, and of SIC during austral summer (December-March) on breeding success of (B) B skuas and (C) NB skuas. SIC effects on breeding success of Adélie penguins during (D) austral summer, (E) austral spring and (F) austral winter (April-Aug). Solid line represents the modeled relationship obtained from the covariate model and shaded areas represent the 95% credibility intervals. Points represent demographic parameter estimates each year plotted against covariate values, with the standard deviations as errors bars. Demographic parameters of south polar skuas depend on their breeding state at the previous breeding season (B: breeder, and NB: non-breeder).

## DISCUSSION

Our results suggest that the dynamics of a seabird predator-prey system was mostly impacted by environmental conditions and driven by bottom-up processes. We showed that the availability of preys played an important role in the survival and breeding parameters of the apex predator. We did not find top-down effects in this system (i.e. effects of skuas on Adélie penguin demographic parameters). Local weather and sea ice conditions during the breeding season and wintering also appeared to be important drivers of breeding success in Adélie penguins and south polar skuas. Important changes of those factors due to global change and their repercussions through bottom-up processes that we observed in this predator-prey system may therefore have major incidence on the future dynamics of both Adélie penguins and south polar skuas.

### Effect of the breeding state of the previous year

As predicted, our results showed differences in estimates and relationships between demographic parameters of skuas, environmental factors and prey-predator relationships as a function of their breeding state during the previous year (see Table 3 and Figure 5), which could reflect different foraging strategies and behaviours.

Both B and NB skuas showed similar demographic responses (survival, breeding success and probability to fledge two chicks) to air temperature and sea ice concentration during spring and summer. This suggests that these environmental parameters similarly affected adult individuals regardless of their previous breeding status. However, NB skuas seemed to be more sensitive to the conditions during wintering than B skuas, as their survival was negatively affected by higher sea temperature anomalies. During the study period there was an increase in sea surface temperature anomalies in the east of Japan where skuas spent the nonbreeding season. High SSTa generally reflect lower availability of food for seabirds through bottom-up effects (Frederiksen et al., 2006; Barbraud et al., 2012; Weimerskirch et al. 2015; Hazen et al., 2019). We therefore suspect an increase in mortality due to decreased food abundance during wintering when sea surface temperature is warmer. NB skuas may be more sensitive to such environmental conditions than B skuas since they might be mostly individuals in poorer body condition, or with less experience. On the opposite, B skuas demographic parameters (survival, breeding probability and probability to fledge two chicks) seemed to be more sensitive to prey availability (abundance of Adélie penguins and number of dead chicks of emperor penguin) than NB skuas. This could be a consequence of the higher energetic costs due to chick rearing compared to NB skuas (Drent & Daan, 1980; Weimerskirch, 1990; Weimerskirch & Lys, 2000).

B skuas had higher breeding probability, higher breeding success, higher probability to fledge two chicks, but lower survival probability than NB skuas. This could indicate the existence of a survival cost of reproduction for B skuas. The large differences between NB and B skuas breeding probability could be a consequence of the limited number of high quality sites. As skuas are faithful to their breeding sites, it might be more difficult for a NB skua to find a place to breed than for a B skua (Young, 1972; Pacoureau et al., 2018). Thus, all individuals cannot breed every year. Less experienced and often less competitive NB skuas could be relegated to poorer qualities sites, more exposed to wind, to snow accumulation and runoff from melting ice and snow (Dhondt et al., 1992; Ferrer & Donazar, 1996; Krüger & Lindström, 2001; McPeek et al., 2001; Kokko et al., 2004). This hypothesis is supported by the lower breeding success in NB skuas compared to B skuas, but must be also be interpreted cautiously since we did not find any effect of density dependence on the breeding probability of NB skuas.

### Density dependence

Density dependence is a well-known factor influencing bird population dynamics (Ashmole, 1963; Fowler, 1981; Turchin, 1995; Newton, 1998). More individuals breeding at the same locality increase intraspecific competition for resources (Charnov et al., 1976), for breeding sites, territorial behaviour (Rodenhouse et al., 1997), and the transmission of diseases or parasites (Coulson, 2002). Several studies have shown negative effects of intraspecific density dependence on long-lived upper trophic bird species showing territorial behaviour like the south polar skuas (Fernandez et al., 1998; Phillips et al., 2004; Turrin & Watts, 2014; Furness, 2015; Quéroué et al., 2021).

We did not detect density dependence effects of the number of skuas on the demographic parameters of NB skuas. However, contrary to our prediction, we found a strong positive effect of the number of adult skuas on the survival of B skuas. Likewise, we also observed a very strong effect of the number of Adélie breeding pairs on the breeding success of Adélie penguins. Those positive relationships are likely an indirect effect of environmental conditions on demographic parameters. When trophic conditions are good, it may favour the number of individuals that come back to the colonies to reproduce, as well as their survival and breeding success.

### Interspecific relationships

The number of Adélie penguin breeding pairs, used as a proxy of the food available for skuas (i.e. Adélie penguins eggs and chicks), had a positive effect on the survival of B skuas (see Figure 6). More Adélie penguins breeding likely increased the number of eggs and chicks available for skuas, which could minimize foraging effort and thus maximize survival for breeding individuals. Therefore, the increase in the number of Adélie penguin breeding pairs that we observed during this study at Pointe Géologie could partly explain the increase in the apparent survival of B skuas.

Contrary to our predictions, we did not find an effect of the number of Adélie penguin breeding pairs on the breeding success of skuas. Although Adélie penguin breeding pairs could be abundant early in the season (Nov-Jan), the extent and timing of egg failure and chick mortality may not match with the most energy demanding period for skuas which occurs during chick rearing (Dec-Mar). This may explain this lack of relationship between skuas breeding performance and numbers of Adélie penguin breeding pairs. Interestingly, we found a positive effect of the number of emperor penguin dead chicks on the probability to fledge two chicks in B skuas (see Figure 6). Mostly consumed early in spring, before eggs and chicks of Adélie penguins become available (Pacoureau et al., 2018), emperor penguin dead chicks will favour skuas to attain a good body condition to afford the energetic costs due to the rearing of their chicks (Williams, 1966; Drent & Daan, 1980; Grilli et al., 2018), and even more to raise two chicks. Thus, food availability early in the breeding season is likely a determinant of the breeding success of skuas.

However, unexpectedly, we observed a negative effect of the number of emperor penguin dead chicks on the breeding probability of B skuas. More food available early in the breeding season should allow more individuals to reach an adequate body condition to breed (Guinet et al., 1998; Madsen & Shine, 1999; Toügo et al., 2002; Reed et al., 2004). We suspect this effect to be indirectly linked unmeasured environmental conditions which could have a negative effect both on the mortality of emperor penguin chicks and on the breeding probability of B skuas. For example, important fast ice extent during austral winter and early spring has a negative effect on the breeding success of emperor penguins (Massom et al., 2009; Labrousse et al., 2021), and may also negatively affect the breeding probability of skuas.

We did not detect a top down effect in this predator-prey system, i.e. a negative effect of the number of adult skuas on the breeding success of Adélie penguins. As the Adélie penguin breeding population is around 340 times larger than the skua breeding population, the predation of skuas on Adélie penguin eggs and chicks is likely negligible.

### Environmental conditions

Local weather conditions during the breeding season influence the breeding success of seabirds directly *via* their effects on offspring growth (Busser et al. 2004; Hahn et al., 2007), and indirectly *via* the body condition of the parents (Grilli et al., 2018). In line with this, we found that this prey-predator system and the breeding success of both skuas and Adélie penguins was highly sensitive to local environmental conditions during the breeding season. To refine our results, it could be interesting in future studies to consider nonlinear relationships when assessing the effects of the covariates.

#### Air temperature

Warmer air temperature in spring had a positive effect on the survival and breeding success of skuas, and on the probability to fledge two chicks in B skuas. Colder air temperature likely increases energetic costs and water loss due to thermoregulation and so decreases body condition of the parents and their chicks (Spellerberg 1969; Pacoureau et al., 2019). Individuals in poorer conditions may lay smaller eggs and of lesser quality with implications on chick quality and mortality until fledging (Furness, 1983; Sæther et al., 1997; Ratclife et al. 1998; Phillips et al. 2004; Hahn et al. 2007).

#### Sea ice concentration

The Adélie penguin is an ice obligate species (Croxall et al. 2002; Ainley, 2002; Jenouvrier et al., 2006) and mainly feed on Antarctic silverfish and krill (Cherel 2008). Thus, as we predicted, Adélie penguin breeding success was strongly driven by sea ice conditions in winter, spring and summer. High SIC in summer had a negative impact on the breeding success of Adélie penguins, as we hypothesised. Extreme sea ice cover during this period increases the distance between the colonies and the foraging areas, which reduces meal frequency for the chicks and may cause abandonment of the nest and chick mortality (Gaston et al. 2005; Ropert-Coudert et al., 2014; Barbraud et al., 2015). In line with this, we observed that the breeding success of Adélie penguins tended to decrease since 1988, which could be partly due to the increasing frequency of extreme summer sea ice concentration events and to an increase of the mean SIC in summer at Pointe Géologie (Figure 3). The negative effect of important SIC in summer (Figure 7) seems to be most likely the result of summers with large expanses of fast ice (correlated to SIC), which are well known to have strong negative effects on breeding success of Adélie penguins (Ropert-Coudert et al. 2014, Barbraud et al. 2015). Decreased winter sea ice is associated with lower densities of krill during summer (Siegel & Loeb, 1995; Loeb et al., 1997), one of the key resources for Adélie penguins during the breeding season. Furthermore, Emmerson & Southwell (2011) showed that the survival of sub-adult and adult penguins was higher when there was high sea ice concentration in winter. Contrary to our prediction, high SIC before the breeding season (in winter and spring) appeared to have a negative impact on the breeding success of Adélie penguins. Tracking studies showed that Adélie penguins preferred foraging habitats correspond to SIC of 20-30% (Le Guen et al., 2018). Medium sea ice cover seems to offer enhanced food availability, as well as reducing the risk of predation (Langbehn & Varpe, 2017). During years when winter and spring SIC are higher, Adélie penguins may be force to move farther away from the colony to find more favourable conditions for foraging. Thus, we can hypothesize that high SIC in winter and spring forces Adélie penguins to draw from their reserves in order to reach more favourable foraging areas, which may influence their body condition and contribute to decrease their breeding success.

Sea ice conditions also had an important effect on the demography of south polar skuas. High SIC in spring favoured survival of NB skuas. This suggests that when high SIC conditions occurred in spring more food resources were available. Higher food resources in spring may also favour NB skuas survival through an increase in their body condition as these individuals are negatively impacted by wintering conditions. Unexpectedly we found a negative effect of high SIC in summer on the breeding success of skuas. Indeed, if high SIC summers coincide with increased mortality of Adélie penguin egg and chicks, one might expect more food available for skuas and a better breeding success. However, as observed in Adélie penguins, the breeding success of skuas decreased since 1988 while mean SIC in summer increased. Moreover, skuas experienced their lower breeding success during years with extreme sea ice concentration (Barbraud et al., 2015). We thus suspect that due to massive dying of Adélie penguin chicks early in the breeding season as it has been observed during years with high summer SIC, skuas might not be able to correctly feed their chicks until fledging because of a lack of available food resources at the end of the rearing period.

## CONCLUSION

We provided here one of the most complete study on the factors impacting the demography of a predator-prey system, the Adélie penguin and south polar skua, while explicitly integrating interspecific relationships. We also proposed a new application of a multispecies IPM for a predator-prey system, in a context where only count time series data are available for one of the species. Estimating demographic parameters and abundance for both species simultaneously while assessing the effects of density-dependence, climatic conditions, and intraspecific relationship in a unique model allows a better propagation of all sources of uncertainty, as well as helping us to better understand joint population dynamics in predator-prey systems in context of climate change. We highlighted that the dynamics of this predator-prey system over the past three decades was mostly driven by bottom-up processes and local environmental conditions. Our results reveal the complexity of the effects of environmental conditions on the demography of a predator-prey system, with several climate and oceanographic variables involved in addition to several interspecific relationships. Overall, high sea ice concentration in spring and in winter, as well as high air temperature in spring had positive effects on south polar skuas, while high sea ice concentration in summer had negative effects on both the predator and its prey. High sea ice concentration in spring and winter also had negative effects of the breeding success of Adélie penguins. This reveals the importance of sea ice before the breeding season for this predator-prey system, which is likely linked to the abundance of marine resources such as krill and Antarctic silverfish, while high sea ice concentration in summer likely increased costs of reproduction due to more constraining foraging conditions. During the past 30 years, populations of both the predator and its main prey have increased, suggesting that the overall net effect of environmental change experienced during the breeding and wintering periods was favourable to these species at this study site. As no negative density dependence was detected, both populations might continue to increase in the future years if climatic conditions remain similar. However, Antarctic climate and oceanography are predicted to suffer important changes by the end of the century (Kirtman et al., 2013), which could have negative impacts on the Antarctic ecosystem and top predators such as seabirds (Jenouvrier et al., 2014).

## ACKNOWLEDGEMENT

The authors thank all the fieldworkers who participated in the long-term studies since 1963 as part of the program IPEV 109 “Seabirds and marine mammals as sentinels of global changes in the Southern Ocean” (PI C. Barbraud). The authors thank D. Joubert for data management. Data were collected with the logistical and financial support from Institut Polaire Français Paul-Emile Victor (IPF), Terres Australes et Antarctiques Françaises and Zone Atelier Antarctique et Terres Australes (LTSER France). This research was funded by the French National Research Agency (grant ANR-16-CE02-0007 DEMOCOM, PI O. Gimenez). C. Barbraud acknowledges support from the BNP Paribas Foundation as part of program SENSEI (SENtinels of the Sea Ice).

## DATA AVAILABILITY STATEMENT

The code and data are available in github (https://github.com/ViollatLise/MultispeciesIPM_skua_adelie.git)

