## Supplementary material for "BOTTOM-UP EFFECTS AND DENSITY DEPENDENCE DRIVE THE DYNAMIC OF AN ANTARCTIC SEABIRD PREDATOR-PREY SYSTEM"

### APPENDIX 1: Details of the integrated population model

Our preys-predator system IPM allows estimating abundance and demographic rates of south polar skuas and Adélie penguins, using capture-recapture and count data for south polar skuas and only times series for Adélie penguins. We built this two species IPM by connecting an IPM for the skuas and a state-space model for Adélie penguins through explicit predator-prey relationships. The two species-specific models used are structured according to life-history traits. We incorporated the effects of species-specific demographic parameters such as apparent survival and breeding parameters like breeding probability and breeding success. We used Poisson (Po) and binomial (Bin) distributions to consider demographic stochasticity.

**Integrated population model for skuas**

The model used here is derived from a model developed for Brown skuas (*Catharacta lönnbergi*) by Quéroué et al. (2021).

***State process***

*Offspring of juvenile skuas*: The number of juveniles aged 0 to 1 years ($N_{J1}$), i.e. the number of chicks produced by successful breeders ($N_{SB1}$ et $N_{SB2}$) according to their fecundity ($f_{SB1} = 1$, $f_{SB2} = 2$, with a sex ratio of 0.5) at the year t, was estimated by a Poisson distribution

$$\mathbf{N_{J1,t} \sim \textit{Po}\ (0.5 \times f_{SB1} \times N_{SB1} + 0.5 \times f_{SB2} \times N_{SB2})} \qquad (1)$$

*Juvenile survival*: The number of juveniles aged between 1 and 2 years ($N_{J2}$), between 2 and 3 years ($N_{J3}$) and between 3 and 4 years ($N_{J4}$) was modeled with a binomial distribution

$$\mathbf{N_{Ji,t} \sim \textit{Bin}\ (\mu_{Ji}, N_{Jx,t-1})} \qquad (2)$$

with i the age of the juvenile state (from $N_{J2}$ to $N_{J4}$), $\mu_{Ji}$ the apparent survival between 1 and 2 years ($\mu_{J1}$), 2 and 3 years ($\mu_{J2}$), and between 3 and 4 years ($\mu_{J3}$). Having no information available to estimate skua survival during their juvenile phase, we assumed that the apparent survival increases linearly with age, the birds becoming more efficient to foraging food or to avoid predators as they acquire experience (Greig et al., 1983; Ainley et al., 1990; Daunt et al., 2007; Grande et al., 2009; Fay et al., 2015). We then have

$$\mathbf{logit\ (\mu_{Ji}) = \lambda_1+\lambda_2 i} \qquad (3)$$

with $\mu_{Ji}$ the apparent survival, i the age of the juvenile state (de $N_{J1}$ à $N_{J4}$), $\lambda_1$ the intercept and $\lambda_2$the slope constrained to be positive

*Juvenile first reproduction*: For skuas the first breeding attempt begins at the age of 4 years old, with a probability Pr. We then have

$$\mathbf{N_{J4B,\,t} \sim \textit{Bin}\ (Pr_{\,t},\ \mu_{J4} \times N_{J4+,\,t-1})} \qquad (4)$$

as the number of individuals aged 4 years old that attempt to breed and

$$\mathbf{N_{J4NB,\,t} = \mu_{J4} \times N_{J4+,\,t-1} - N_{J4B,\,t},} \qquad (5)$$

as the number of individuals aged 4 years old that don't attempt to breed, with $\mu_{J4}$ the apparent survival for the state $N_{J4+}$. $N_{J4+}$ included individuals in their 4 years ($N_{J4}$) and 4 years old or older individuals that did not attempt to breed during previous breeding seasons ($N_{J4NB}$).

*Adult survival*: We modeled the number of surviving adults at the year t among the total number of adults at the year t-1 with µ an apparent survival of adult and a binomial distribution

$$\mathbf{N_{alive,\,t} \sim \textit{Bin}\ (\mu_{\,t-1},\ N_{adtot,\,t-1})} \qquad (6)$$

with $N_{adtot,\,t}$ the sum of breeder ($N_{B,\,t}$) and non-breeder ($N_{NB,\,t}$) individuals at the year t, calculated as

$$\mathbf{N_{adtot,\,t} = N_{B,\,t} + N_{NB,\,t}} \qquad (7)$$

*Breeding probability*: The number of individuals having reproduced or not among individuals that survived is modeled with a binomial distribution as

$$\mathbf{N_{B\ alive,\,t} \sim \textit{Bin}\ (\beta_{\,t},\ N_{alive,\,t})} \qquad (8)$$

$$\mathbf{N_{NB,\,t} = N_{alive,\,t} - N_{B\ alive,\,t}} \qquad (9)$$

with β the breeding probability, $N_{B\ alive,\,t}$ the breeders that survived, and $N_{NB,\,t}$ the non-breeders that survived. The total number of breeders that survived was defined by breeders that survive ($N_{B\ alive}$), the number of juveniles that attempted to breed for the first time ($N_{J4B}$), and immigrants ($N_{im}$) i.e. new individuals observed for the first time at the colony

$$\mathbf{N_{B,\,t} = N_{B\ alive,\,t} + N_{J4B,\,t} + N_{im,\,t}} \qquad (10)$$

*Breeding success*: The number of individuals that were failed breeders ($N_{FB}$) and individuals that have succeeded in their breeding ($N_{SB}$) were modeled following a binomial distribution with ϕ the probability of breeding success.

$$\mathbf{N_{SB,\,t} \sim \textit{Bin}\ (\phi_{t-1},\ N_{B,\,t})} \qquad (11)$$

$$\mathbf{N_{FB,\,t} = N_{B,\,t} - N_{SB,\,t}} \qquad (12)$$

A successful breeder can fledge one chick ($N_{SB1}$) or two chicks ($N_{SB2}$), and this process was modeled with a binomial distribution and δ the probability of successfully fledging 2 chicks

$\mathbf{N_{SB2,\,t} \sim \textit{Bin}\ (\delta_{t-1},\ N_{SB,\,t})}$ (13)

$\mathbf{N_{SB1,\,t} = N_{SB,\,t} - N_{SB2,\,t}}$ (14)

***Observation process***

*Count data*: An observation equation allows to link the number of breeding pairs of skuas estimated by the state process ($N_B$) with the number of breeding pairs observed (Ys) at year t, while considering $\varepsilon_t$ an observation error, and following a normal distribution

$\mathbf{Ys_t \sim \textit{N}\ (N_{B,\,t},\ \varepsilon_t)}$ (15)

with $\boldsymbol{\varepsilon_t \sim \textit{N}\ (0,\ \sigma^2_{Ys})}$ the error term and its variance. We thus have the following likelihood for the count data for skua:

$\mathbf{L_{CO}\ (Ys \mid \mu_{J1},\ \mu_{J2},\ \mu_{J3},\ \mu_{J4},\ Pr,\ \mu,\ \beta,\ \gamma,\ \delta,\ N_B)}$ (16)

*Capture-recapture data*: To estimate the demographics parameters of skuas from the CR data, we used a multi-state capture-recapture model (Kéry & Schaub, 2012). This model involved five states: NB, FB, SB1, SB2, D (dead) and six events: not observed, observed as NB, observed as FB, observed as SB1, observed as SB2, and observed but with an uncertain state. The following demographic parameters were estimated: (μ) the apparent survival probability, (β) the breeding probability, (ϕ) the breeding success, and (δ) the breeding success with 2 chicks. This model allowed considering imperfect detection of individuals and uncertainty during the state assignment (Gimenez et al., 2012). We thus estimated two additional parameters: the detectability probability (p) and the probability to assign an individual to a state (c). All the parameters, excluding (c), included a yearly random temporal effect.

The transition of an individual from a given state during the breeding season t depended on its breeding states at the previous breeding season t-1: breeders (B) represented individuals that have bred during the previous season (individuals in states FB, SB1, or SB2) and nonbreeders (NB) represented individuals that did not attempt to breed at the previous season. Thus, demographic parameters of an individual were dependent on its breeding states at the previous breeding season.

$\mathbf{L_{CR}\ (CH \mid \mu,\ \beta,\ \phi,\ \delta,\ p,\ c)}$ (17)

*Joint likelihood*: The joint likelihood for the skua IPM was the product of the likelihood for the count data ($L_{CO}$) and the likelihood for the capture-recapture data ($L_{CR}$)

$\mathbf{L_{IPM}\ (Ys,\ CH \mid \mu_{J1},\ \mu_{J2},\ \mu_{J3},\ \mu_{J4},\ Pr,\ \mu,\ \beta,\ \gamma,\ \delta,\ N_B,\ p,\ c)}$

$\mathbf{= L_{CO}\ (Ys \mid \mu\ J1,\ \mu\ J2,\ \mu\ J3,\ \mu\ J4,\ Pr,\ \mu,\ \beta,\ \gamma,\ \delta,\ N_B) + LCR\ (CH \mid \mu,\ \beta,\ \phi,\ \delta,\ p,\ c)}$ (18)

**State space model for Adélie penguins**

Since we did not have any data about non-breeders in the population, we built the model on the breeding part of the population of Adélie penguins at Pointe Géologie

***State process***

*Adult survival*: We modeled the number of breeding pairs surviving at year t among the number of breeders at the year t-1 considering an apparent survival probability $\mu_A$, and following a binomial distribution

$$N_{alive,t} \sim Bin(\mu_{A,t}, N_{AB,t-1}) \quad (19)$$

Number of breeding *adults*: We estimated that all alive individuals at year t ($N_{alive}$) breed. Every year, new breeders ($N_{NR}$) arrived at the colony, either from immigration, or local recruits for their first reproduction. We fixed a recruitment rate φ, corresponding to 30% of the breeding population, according to the results of Hinke et al. (2014). The number of new breeders ($N_{NR}$) and the total number of breeders ($N_B$) are thus calculated as

$$N_{NR,t} = \varphi \times N_{alive,t} \quad (20)$$

$$N_{AB,t} = N_{alive,t} + N_{NR,t}. \quad (21)$$

*Breeding success*: The number of fledge chicks (P) born during the breeding season t was estimated following a Poisson distribution, with $\phi_A$ the breeding success

$$P_t \sim Po(N_{AB,t} \times \phi_A) \quad (22)$$

**Observation process**

*Count data*: Two equations allowed linking the number of Adélie penguin breeding pairs ($N_B$) and the number of fledge chicks (P) estimated by the model with the observed number of breeding pairs ($Y_A$), and with the observed number of fledge chicks ($Y_P$) at year t, considering an observation error εt, with εt ~ N (0, $\sigma^2 y_A$), ε the error term and $\sigma^2 y$ its variance, following a normal distribution

$$Y_{A,t} \sim N(N_{AB,t}, \varepsilon_t) \quad (23)$$

$$Y_{P,t} \sim N(P_t, \varepsilon_t) \quad (24)$$

*Likelihood*: We thus have the following likelihoods ($L_A$) for the count data of Adélie penguin breeding pairs and ($L_P$) for fledge chicks

$$L_A(Y_A \mid \mu_A, \varphi, N_{AB}) \quad (25)$$

$$L_A(Y_P \mid \phi_A, P) \quad (26)$$

### APPENDIX 2: ASSESSING THE EFFECT OF ENVIRONMENTAL COVARIATES AND POPULATION DENSITIES

To estimate the effect of environmental covariates (T°, SIC, SSTa) as well as intra and interspecific relationships on the demographic parameters of skuas and Adélie penguins, we used logit linear regressions inside our multispecies IPM.

For example, we modeled the survival ($\mu_B$) of breeding skuas using a logit link:

$$\mathbf{logit\ (\mu B) = \mathit{a} + \alpha N_{AB,t} \times N_{AB,t} + \alpha N_{adtot,t} \times N_{adtot,t} + \alpha N_{emp} \times N_{emp,t} + \alpha T°_{SN,t} \times T°_{SN,t} +}$$

$$\mathbf{\alpha T°_{DM,t} \times T°_{DM,t} + \alpha SIC_{SN,t} \times SIC_{SN,t} + \alpha SIC_{DM,t} \times SIC_{DM,t} + \alpha SSTa_{t} \times SSTa_{t} + \varepsilon_{\mu B,\, t}}$$

$$\mathbf{\varepsilon_{\mu B,\, t} \sim Norm(0, \sigma^2_{\varepsilon \mu B})}$$

with $a$ the intercept, $\alpha N_{AB}$ the slope indicating the strength of the predator-prey relationship between skuas and Adélie penguins with $N_{AB}$ the number of Adélie penguin breeding pairs, $\alpha N_{adtot}$ the slope indicating the strength of the density dependence with $N_{adtot}$ the number of adult skuas (breeders and non-breeders), $\alpha N_{emp}$ the slope indicating the strength of the predator-prey relationship between emperor penguins and skuas with $N_{emp}$ the number of emperor penguin dead chicks, $\alpha T°_{SN}$ the slope for the effect of air temperature in spring (September-November), $\alpha T°_{DM}$ the slope for the effect of air temperature in summer (December-March), $\alpha SIC_{SN}$ the slope for the effect of SIC in spring, $\alpha SIC_{DM}$ the slope for the effect of SIC in summer and $\alpha SSTa$ the slope for the effect of SSTa in wintering season. $\varepsilon_{\mu B}$ is a yearly random temporal effect and $\sigma^2_{\varepsilon \mu B}$ its temporal variance.

### APPENDIX 3: Estimation of the capture probability for south polar skuas

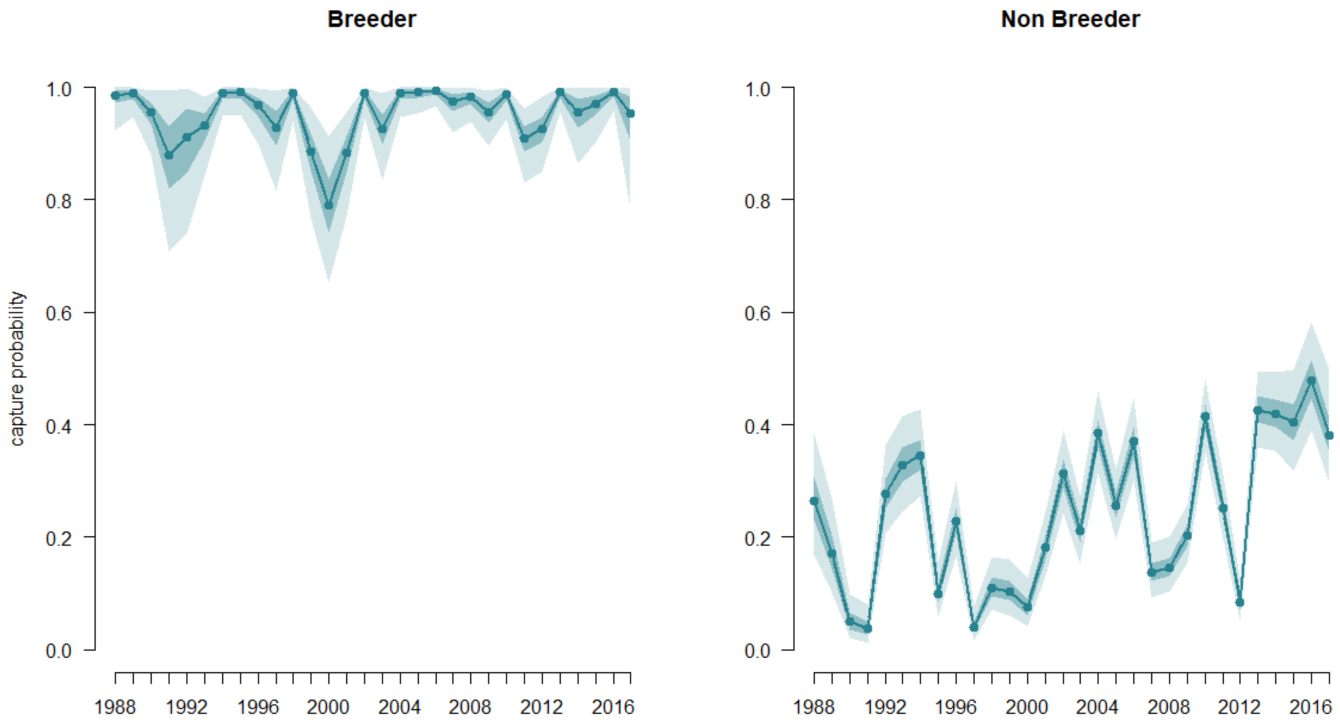


**FIGURE A1.1** Estimated capture probability for south polar skuas of Pointe Géologie from 1988 to 2018, depending on their breeding state at the previous breeding season. Solid lines represent the means of marginal posterior distributions. Shaded area are the 50% and 95% credibility intervals.

**TABLE A1.1** Capture probability for south polar skuas depending on their breeding state (B: breeder and NB: non-breeder) at the previous breeding season estimated by the model from 1988 to 2018 for the populations of Pointe Géologie, with the 95 % confidence interval (CRI)

| **Parameter** | | **Mean** | **CRI** |
|---|---|---|---|
| **Capture probability** | B | 0.945 | (0.867, 0.991) |
| | NB | 0.242 | (0.177, 0.32) |

### APPENDIX 4: Estimation and variation of the abundances of the different breeding states of south polar skuas

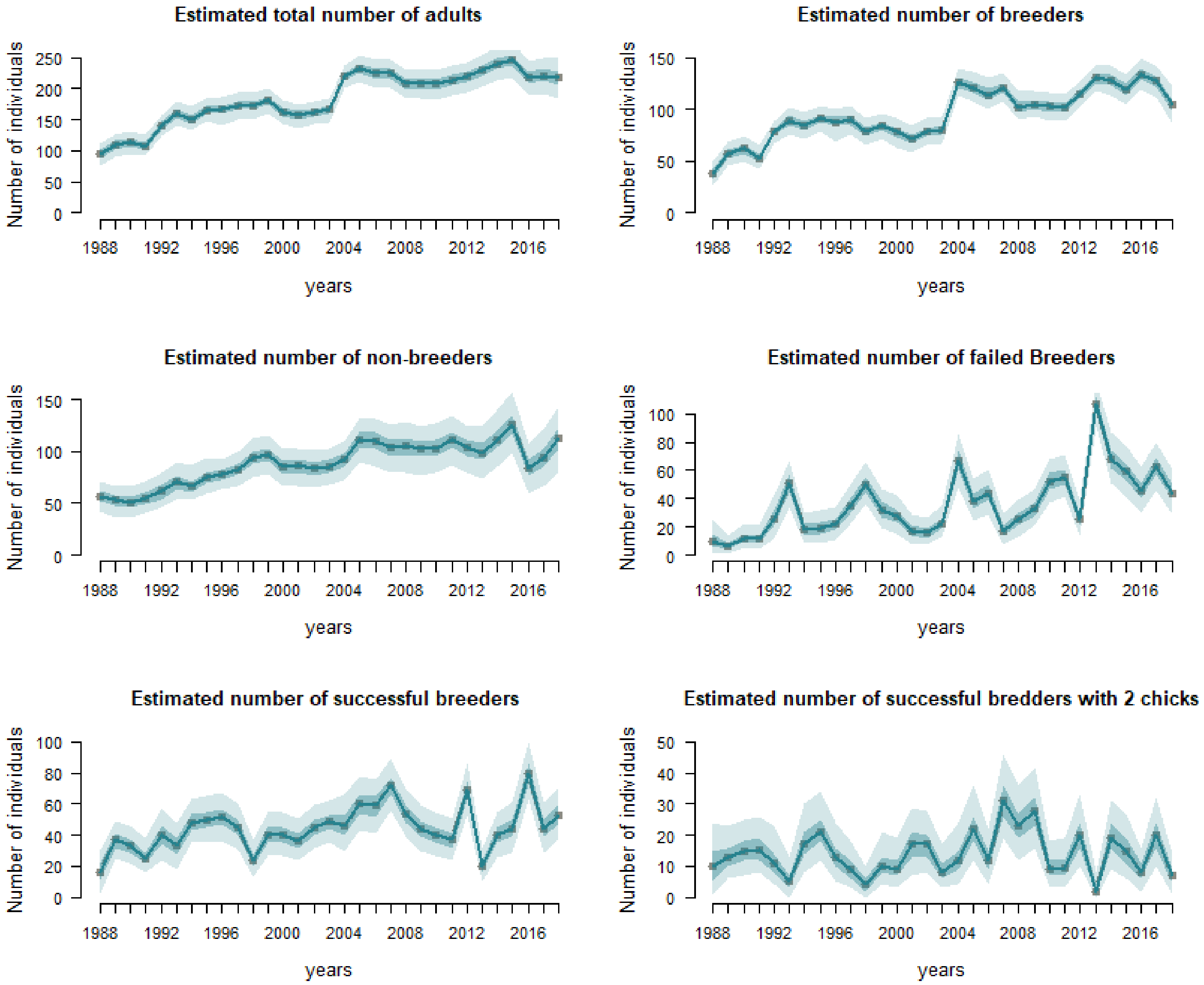


**FIGURE A2.1** Estimated number of individuals from different adult breeding states for each year (1988-2018) for south polar skuas. Solid lines represent the means of marginal posterior distributions. Shaded area are the 50% and 95% credibility intervals.

### APPENDIX 5: Estimation of the effects of environmental covariates, prey-predator relationships and intraspecific density-dependence on the demographic parameters of south polar skuas and Adélie penguins

**TABLE A3.1**: Estimation of the effects of environmental covariates, prey-predator relationships and intraspecific density dependence on the demographic parameters of south polar skuas depending on their state at the previous breeding season (breeder or non-breeder), with the mean slope of the effect estimated by the model, the 95 % confidence interval (CRI) and the probability to have a slope with a negative value (PN). The effect considered as significant, i.e. estimated effect with a probability of being negative (PN) greater than 90% or less than 10%, are in bold.

| | Adult apparent survival μ | | | | | |
|---|---|---|---|---|---|---|
| | Breeder | | | Non Breeder | | |
| Covariates | slope | CRI | PN | slope | CRI | PN |
| P emperor penguin | 0.036 | ( -0.318, 0.406) | 0.419 | 0.207 | ( -0.681, 0.937) | 0.328 |
| PP Adélie penguin | **0.76** | **( 0.351, 0.993)** | **0** | -0.039 | ( -0.955, 0.94) | 0.519 |
| DD | **0.882** | **( 0.687, 0.996)** | **0** | -0.121 | ( -0.964, 0.913) | 0.596 |
| AT spring | **0.225** | **( -0.08, 0.539)** | **0.066** | **0.435** | **( -0.131, 0.942)** | **0.057** |
| AT summer | 0.126 | ( -0.177, 0.424) | 0.186 | 0.132 | ( -0.608, 0.849) | 0.342 |
| SIC spring | 0.062 | ( -0.217, 0.351) | 0.327 | **0.482** | **( -0.199, 0.956)** | **0.063** |
| SIC summer | 0.179 | ( -0.144, 0.509) | 0.1265 | -0.207 | ( -0.833, 0.415) | 0.774 |
| SSTa | 0.045 | ( -0.238, 0.332) | 0.3825 | **-0.544** | **( -0.972, 0.251)** | **0.931** |

| | Breeding probability β | | | | | |
|---|---|---|---|---|---|---|
| | Breeder | | | Non Breeder | | |
| Covariates | slope | CRI | PN | slope | CRI | PN |
| P emperor penguin | **-0.206** | **( -0.458, 0.016)** | **0.966** | -0.036 | ( -0.253, 0.175) | 0.637 |
| PP Adélie penguin | 0.004 | ( -0.925, 0.911) | 0.499 | 0.421 | ( -0.684, 0.976) | 0.184 |
| DD | -0.039 | ( -0.807, 0.827) | 0.542 | 0.151 | ( -0.664, 0.837) | 0.336 |
| AT spring | -0.068 | ( -0.308, 0.168) | 0.705 | -0.011 | ( -0.234, 0.207) | 0.539 |
| SIC spring | 0.17 | ( -0.112, 0.445) | 0.11 | -0.13 | ( -0.366, 0.108) | 0.852 |
| SSTa | -0.106 | ( -0.355, 0.132) | 0.796 | -0.094 | ( -0.321, 0.128) | 0.814 |

| | Breeding success ϕ | | | | | |
|---|---|---|---|---|---|---|
| | Breeder | | | Non Breeder | | |
| Covariates | slope | CRI | PN | slope | CRI | PN |
| P emperor penguin | 0.192 | ( -0.207, 0.607) | 0.18 | 0.263 | ( -0.174, 0.682) | 0.108 |
| PP Adélie penguin | -0.161 | ( -0.968, 0.91) | 0.6 | -0.177 | ( -0.963, 0.919) | 0.642 |
| DD | -0.224 | ( -0.963, 0.878) | 0.712 | -0.424 | ( -0.973, 0.573) | 0.834 |
| AT spring | **0.251** | **( -0.168, 0.615)** | **0.095** | **0.289** | **( -0.085, 0.691)** | **0.071** |
| AT summer | -0.09 | ( -0.472, 0.304) | 0.676 | -0.268 | ( -0.683, 0.203) | 0.896 |
| SIC spring | 0.002 | ( -0.364, 0.357) | 0.484 | -0.172 | ( -0.574, 0.268) | 0.816 |
| SIC summer | **-0.462** | **( -0.882, -0.044)** | **0.986** | **-0.525** | **( -0.916, -0.121)** | **0.994** |

| | Breeding success with two chicks δ | | | | | |
|---|---|---|---|---|---|---|
| | Breeder | | | Non Breeder | | |
| Covariates | slope | CRI | PN | slope | CRI | PN |
| P emperor penguin | **0.309** | **( -0.038, 0.695)** | **0.039** | -0.233 | ( -0.877, 0.448) | 0.751 |
| PP Adélie penguin | -0.283 | ( -0.972, 0.804) | 0.712 | -0.013 | ( -0.942, 0.95) | 0.507 |
| DD | -0.056 | ( -0.928, 0.926) | 0.56 | 0.091 | ( -0.939, 0.95) | 0.417 |
| AT spring | **0.262** | **( -0.047, 0.615)** | **0.052** | 0.019 | ( -0.605, 0.571) | 0.448 |
| AT summer | -0.056 | ( -0.376, 0.266) | 0.646 | 0.153 | ( -0.498, 0.784) | 0.32 |
| SIC spring | 0.075 | ( -0.249, 0.404) | 0.314 | -0.06 | ( -0.731, 0.557) | 0.573 |
| SIC summer | -0.216 | ( -0.623, 0.164) | 0.864 | 0.202 | ( -0.476, 0.887) | 0.286 |

***Notation**: AT: air temperature during spring (AT spring) and during summer (AT summer). SIC: sea ice concentration during spring (SIC spring), summer (SIC summer). Spring corresponds to the breeding period when pairs are formed, eggs are laid and incubated (between September and November), summer corresponds to the chick rearing period, after the eggs hatch until chicks fledge (between December and March). SSTa: sea surface temperature anomalies encountered by skuas during their wintering off the coast* *of Japan (May-June). DD: intraspecific density dependence. PP Adélie penguin: number of Adélie penguin breeding pairs. PP emperor penguin: number of emperor penguin dead chicks.*

**TABLE A3.2**: Estimation of the effects of environmental covariates, prey-predator relationships and intraspecific density dependence on the breeding success of Adélie penguins, with the mean slope of the effect estimated by the model, the 95 % confidence interval (CRI) and the probability to have a slope with a negative value (PN). The effect considered as significant, i.e. estimated effect with a probability of being negative (PN) greater than 90% or less than 10%, are in bold.

***Notation****: SIC: sea ice concentration during spring (SIC spring), summer (SIC summer) and winter (SIC winter).*

| | **Adélie penguin breeding success $\phi A$** | | |
|---|---|---|---|
| Covariates | slope | CRI | PN |
| PP skua | 0.013 | (-0.017, 0.035) | 0.169 |
| SIC spring | **-0.643** | **(-0.663, -0.618)** | **1** |
| SIC summer | **-0.999** | **(-1, -0.997)** | **1** |
| SIC winter | **-0.981** | **(-0.999, -0.954)** | **1** |
| DD | **0.999** | **(0.998, 1)** | **0** |

*Spring corresponds to the breeding period when pairs are formed, eggs are laid and incubated (between September and November), summer corresponds to the chick rearing period, after the eggs hatch until chicks fledge (between December and March), and winter corresponds to conditions encountered by Adélie penguins outside the breeding season in the wintering area (between April and august). DD: intraspecific density dependence. PP skuas: number of adult skuas.*
